# Electrostatic interactions compensate energetic frustration in the coupled folding and binding of intrinsically disordered proteins

**DOI:** 10.64898/2026.01.07.698246

**Authors:** Lucía Álvarez, Nicolás A. Garrone, Michael Claron, Kristian Schweimer, Juliana Glavina, David Will, Peter Sehr, Joe Lewis, Toby J. Gibson, Ignacio E. Sanchez, Janosch Hennig, Lucía B. Chemes

**Affiliations:** Instituto de Investigaciones Biotecnológicas, Universidad Nacional de San Martín (UNSAM), Consejo Nacional de Investigaciones científicas y técnicas (CONICET); Escuela de Bio y Nanotecnologías (EByN), Universidad Nacional de San Martín, Av. 25 de Mayo y Francia, Buenos Aires CP 1650, Argentina; Molecular Systems Biology Unit, European Molecular Biology Laboratory, 69117 Heidelberg, Germany; Chemical Biology Core Facility, European Molecular Biology Laboratory, 69117 Heidelberg, Germany; Chair of Biochemistry IV, Biophysical Chemistry, University of Bayreuth, 95447 Bayreuth, Germany; Laboratorio de Fisiología de Proteínas, Instituto de Química Biológica- Facultad de Ciencias Exactas y Naturales IQUIBICEN-Conicet, Universidad de Buenos Aires, (1428), Buenos Aires Argentina

## Abstract

Intrinsically disordered protein regions (IDRs) establish highly specific protein-protein interactions that play key roles in cell signaling. Many IDRs adopt an ordered structure upon binding to folded protein domains, but the energy landscape for coupled folding and binding (CFB) is poorly understood. Here, we elucidate the energy landscape for CFB using the LxCxE motif from the human papillomavirus E7 protein (LxCxE_WT_) as a minimal model system. Kinetic and structural analysis uncovers strong compensatory energetics whereby stabilizing electrostatic interactions counterbalance energetic frustration in the rate-limiting step for CFB. This mechanism, which we refer to as electrostatic compensation, enables a dynamic search for native contacts while preventing complex dissociation. A global analysis reveals energetic frustration in earliest steps of CFB for many IDRs. Electrostatic compensation may be a widespread mechanism evolved to allow fine-tuning of the affinity and specificity of IDR interactions, which dictates functional selection for charge content within IDRs.

## INTRODUCTION

Intrinsically disordered proteins and protein regions (collectively IDRs) make up 40% of the human proteome. IDRs play key roles in signaling pathways by mediating the formation of highly specific and readily switchable protein-protein interactions^1,2^, whose dysregulation is a hallmark of human disease^3,4^. Numerous studies demonstrate that many IDRs involved in cellular signaling form rigid structures when bound to ordered protein domains^1^. These coupled folding and binding (CFB) events often follow an induced-fit mechanism^5^ in which folding occurs following the binding of an IDR to its partner^6–9^. CFB of IDRs is fast and frequently steered by electrostatic interactions^10,11^. A complete understanding of CFB requires characterizing the energy landscapes that IDRs sample when they bind their partner and identifying the bottleneck of the reaction^12–14^. This bottleneck provides key information on the forces that drive the reaction and is defined by the ensemble of structures that populate the rate-limiting barrier for association, also referred to as the transition state ensemble (TSE).

Kinetic studies combined with extensive mutational scanning were used to characterize the energy landscape for folding of small globular domains^15,16^. These minimal model systems revealed that folding TSEs are composed of protein chains that present a collapsed conformation where many native contacts have formed^17–19^. These observations support a funneled energy landscape whereby protein folding is driven mainly by the native topology^20^. In contrast, many CFB reactions appear to have a relatively flat free energy landscape where the TSE is disordered and most native contacts form after the rate-limiting step^9,21–27^. This is consistent with the structurally heterogeneous encounter complexes reported in NMR^7,28^, smFRET^29,30^ and molecular dynamics (MD) simulations^31,32^. MD simulations also predict the existence of transient non-native contacts^31,33^ that must reconfigure to allow the formation of the native complex and would create a more rugged energy landscape. However, limited experimental data^21^ questions whether non-native contacts are present in the TSE for CFB. Moreover, a disordered TSE does not explain which forces reduce the dimensionality of the search for the native state. Non-specific hydrophobic interactions stabilize some CFB TSEs, but they do not explain most systems^27^. Electrostatic interactions mediate the rapid association of IDRs with their binding partners through electrostatic steering^10^, and they could play additional roles to drive the formation of the folded complex.

Short IDRs that undergo CFB^34,35^ provide advantages as minimal model systems to elucidate the energy landscape of CFB reactions. The LxCxE motif from the human papillomavirus 16 E7 protein (LxCxE_WT_) is a short IDR that folds upon binding to the retinoblastoma protein pocket domain (Rb) and it can serve as an ideal model system for CFB for several reasons. First, LxCxE_WT_ CFB occurs via a two-state mechanism that simplifies kinetic analysis^36^ (Figure 1A). Second, extensive mutational scanning is feasible because viral cell-cycle hijack ^37^ led to the evolution of high binding affinity (*K_D_* = 5 nM) for LxCxE_WT_ ^38–41^. Third, unbound LxCxE_WT_ is disordered^42,43^ and it adopts an extended structure devoid of canonical secondary structure upon CFB^44^ (Figure 1B), reducing contributions from ground state effects that confound the analysis of TSE energetics^45,46^.

**Figure 1.**
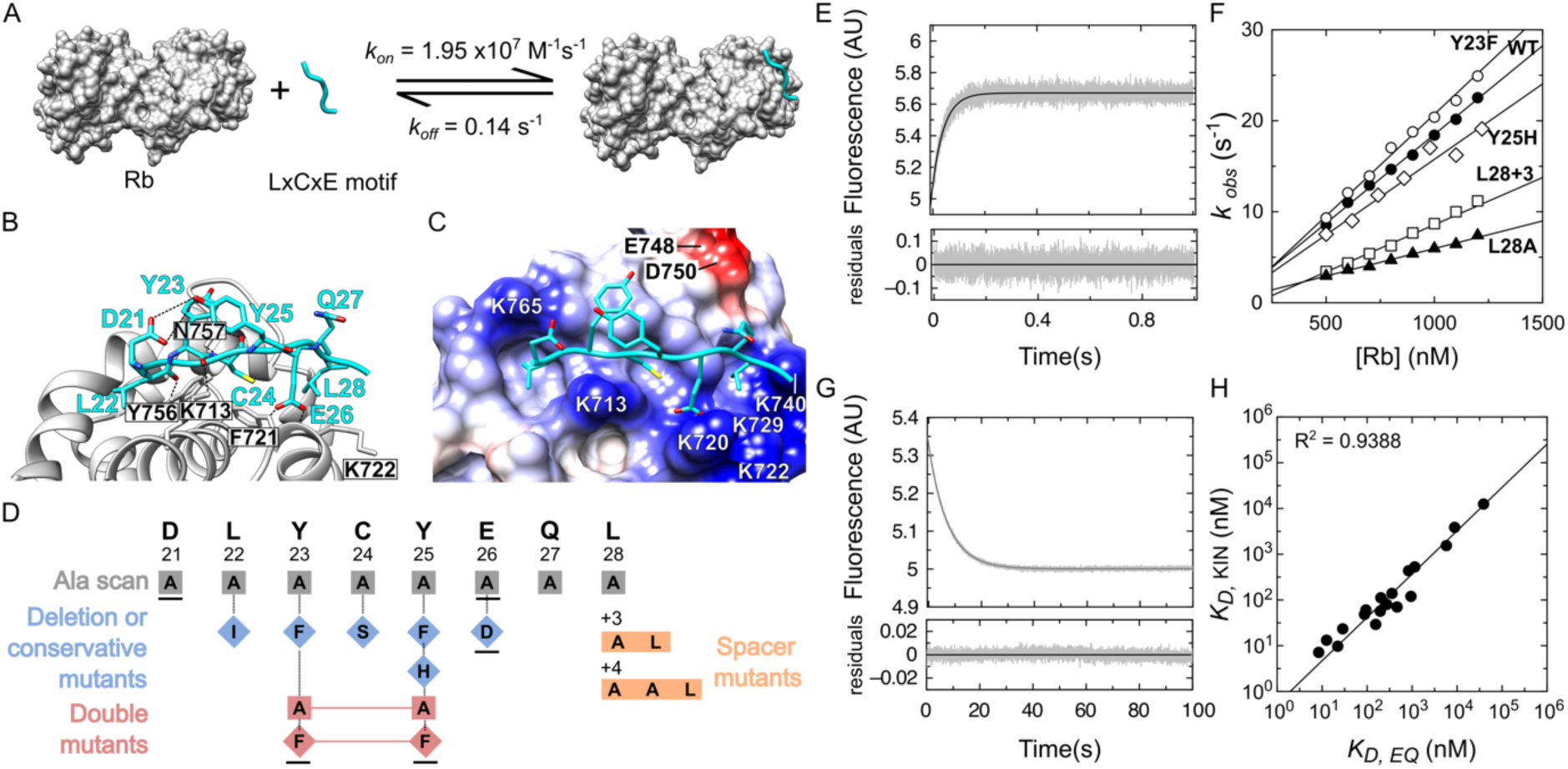
LxCxE_WT_ as a minimal model system for CFB. A) Two-state reaction for CFB of LxCxE_WT_ to Rb (PDB ID 1GUX). B) LxCxE_WT_ (cyan) bound to the Rb LxCxE cleft (grey). C) Rb domain rendered using coulombic surface coloring (blue = positive charge; red = negative charge) highlighting charged residues that line the LxCxE cleft. D) Schematic representation of the mutational strategy used in this study. LxCxE_WT_ sequence and residue numbering are shown at the top. Gray squares: Alanine scan; Blue diamonds: Deletion or Conservative mutations; Orange boxes: Spacer mutants; Pink diamonds: Double mutants. Black underline: Mutants subjected to electrostatic dissection. Vertical dotted lines indicate positions for which relative thermodynamic parameters between different substitutions were derived. E) Association trace following addition of 1.2 μM Rb to 100 nM FITC-LxCxE_WT_. Black line: Fit to a mono-exponential decay (k_obs_ = 22.5 ± 0.1 s^-1^, residuals below the plot). F) PFO plots for selected LxCxE motif mutants. G) Dissociation trace following displacement of a 100nM [FITC-LxCxE_WT_:Rb] complex with 20-fold excess (2 μM) unlabeled LxCxE_WT_. Black line: Fit to a mono-exponential function (k_off_ = 0.139 ± 0.001 s^-1^, residuals below the plot). h) Plot of *K_D, KIN_* versus *K_D,_* _EQ_ for all LxCxE mutants. Black line: linear correlation (slope = 0.87, R^2^ = 0.94). *K_D,_* _EQ_ values are on average 2-fold the *K_D, KIN_* value in agreement with previous reports showing that *K_D,_* _EQ_ values exceed those obtained by direct titration by 2-3 fold^38^. All kinetic traces, PFO plots and *K_D, KIN_/K_D,_* _EQ_ values are reported in SI Appendix, Figures S1-S3 and Tables S2-S3.

Here, we combine kinetic analysis with mutational scanning to elucidate the energy landscape for CFB of LxCxE_WT_ to Rb and we experimentally rule ground state effects using structural analysis. By dissecting the contribution of electrostatic and neutral components we identify energetic frustration created by non-native contacts in the TSE for CFB. Further, we uncover a compensatory mechanism through which electrostatic interactions offset energetic frustration, that we term electrostatic compensation. Electrostatic compensation explains how an early and disordered TSE can overcome the energetic frustration created by non-native contacts and reconfigure to yield the native complex. A global analysis extends the findings with LxCxE_WT_ and identifies energetic frustration in the TSE for CFB of many IDRs, which is associated with residues that stabilize the native complex. This suggests that electrostatic compensation may be a general mechanism evolved to allow fine-tuning of the affinity and specificity of IDR interactions while limiting crosstalk and promiscuous binding.

## RESULTS

### Mutational scanning approach for assessing the CFB of LxCxE_WT_

LxCxE_WT_ folds upon binding to Rb via four contact positions (L22/C24/L28/E26) that establish intermolecular hydrophobic and hydrogen bond interactions with the LxCxE cleft on Rb ^44^. In addition, four solvent-exposed non-contact positions (D21/Y23/Y25/Q27) establish stabilizing intramolecular interactions that include a hydrogen bond between D21 and Y25 and a π-stacking interaction between Y23 and Y25^40,47^ (Figure 1B). Long-range electrostatic interactions between negatively charged LxCxE_WT_ and the positively charged rim of the LxCxE cleft speed up association through electrostatic steering^36^ (Figure 1C), recapitulating the effect of charges in many CFB reactions^11,48^. To assess the contribution of contact and non-contact positions to the energetics of CFB, we performed alanine scanning for residues D21-L28 (Figure 1D). We also introduced structurally conservative mutants representing variants present in other LxCxE motifs (L22I, C24S, E26D), including the variable spacing of the L28 position^47^. Last, we replaced Y23 and Y25 with Phe and Ala to assess the contribution from intramolecular H-bond formation and aromatic ring stacking to the binding energetics. This comprehensive set of mutations (Figure 1D and SI Appendix, Table S1) takes advantage of the sequence space explored by LxCxE motifs, which span a broad range of binding affinities^47,49^.

### LxCxE_WT_ as a minimal model system for CFB

Kinetic analysis of LxCxE_WT_ binding to Rb using stopped flow fluorescence revealed mono-exponential association and dissociation reactions and a linear dependence of *k_obs_* on Rb concentration under pseudo-first order conditions (Figures 1 E-G). The kinetic rate constants (Figure 1A, SI Appendix, Table S2) were in good agreement with the previously reported two-state reaction mechanism^36^. The mutants we designed showed similar two-state behavior to LxCxE_WT_ (SI Appendix, Figures S1-S3, Table S2) and a high correlation between *K_D_* values obtained by kinetic analysis (*K_D,KIN_* = *k_off_/k_on_*) and those derived from equilibrium measurements (*K_D,EQ_*) using an orthogonal high-sensitivity fluorescence plate assay (R^2^ = 0.94) (Figure 1H and SI Appendix, Table S2 and Figure S4). This indicates that all mutants follow a two-state reaction mechanism, allowing a straightforward interpretation of their effects on the TSE energetics.

We first assessed the effect of mutations on global stability and on the rate-limiting barrier for association by calculating the changes in the free energy of binding and activation (*ΔΔG^0^/ΔΔG**^‡^***) for each mutant with respect to LxCxE_WT_ (Eqs. (3-6), Materials and Methods and SI Appendix and Table S2). All mutations were destabilizing at equilibrium (*ΔΔG^0^* >0), indicating that the LxCxE_WT_-Rb complex is optimized for high affinity binding (Figure 2 A, B). Alanine scanning revealed the presence of “hot-spots” for binding (*ΔΔG^0^* > 2 kcal/mol)^50^ at the motif-defining (L/C/E) positions. Mutating the L28 contact position had an effect similar to mutating non-contact positions (D21/Y23/Y25). Conversely, Y23A-Y25A had an effect comparable to mutating a “hot spot” position (Figure 2 A, B). This indicates that while contact positions are determinant for CFB of LxCxE_WT_ to Rb, stabilizing intramolecular interactions from non-contact positions contribute strongly to fine-tuning the binding affinity.

**Figure 2.**
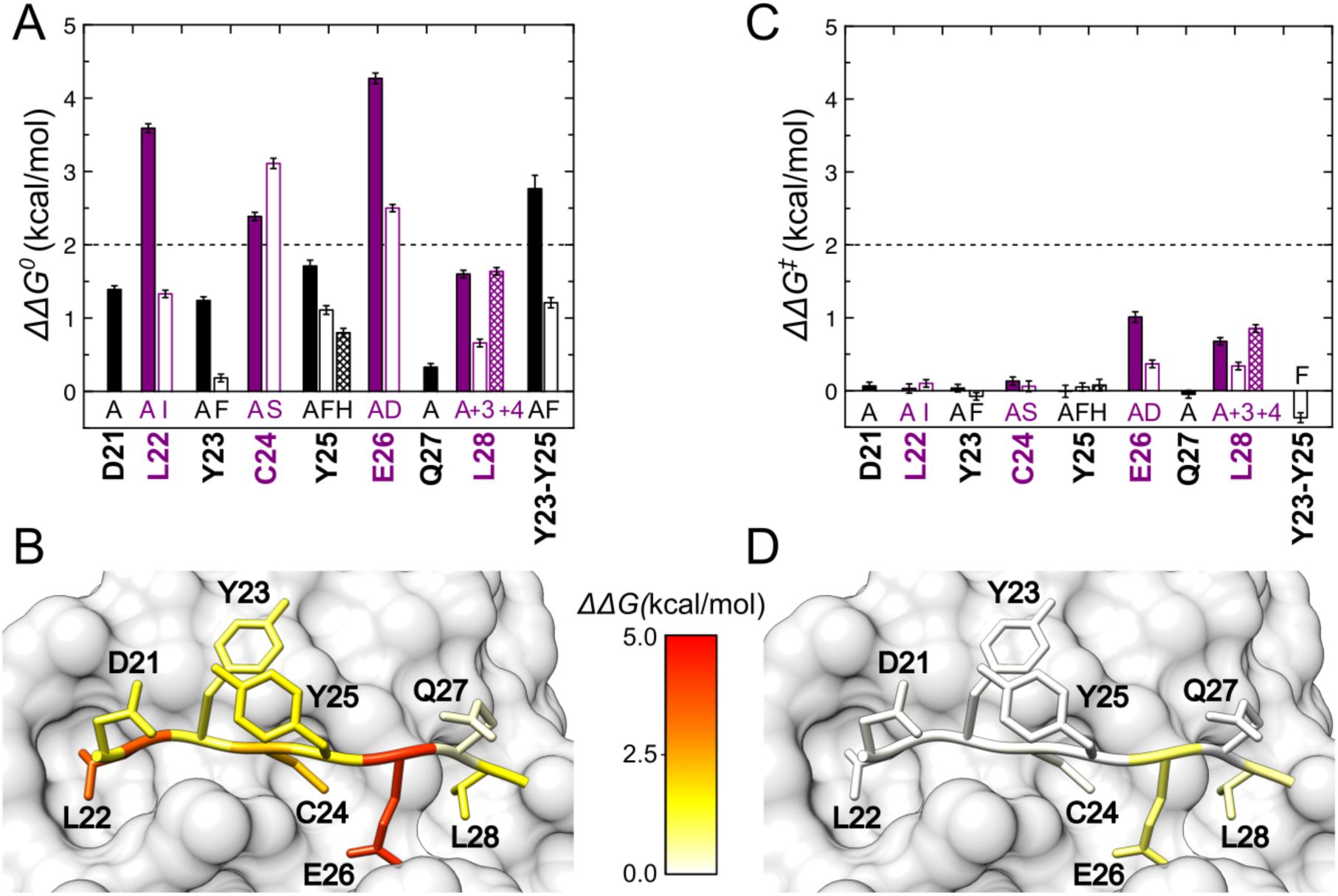
Thermodynamic landscape for LxCxE_WT_ CFB. A) Change in free energy of interaction at equilibrium (*ΔΔG^0^*) for LxCxE motif mutants at 200mM [NaCl]. *ΔΔG^0^* values were obtained from kinetic data except for the Y23A-Y25A mutant, for which *ΔΔG^0^* was calculated from an estimated *K_D,kin_* value obtained from *K_D,comp_* using the empirical relationship established in Figure 1H (SI Appendix, Tables S2). B) Structure of the LxCxE_WT_:Rb complex (PDB ID: 1GUX) colored according to the per-residue ΔΔG° values. C) Change in free energy of interaction for activation (*ΔΔG^‡^*) for LxCxE motif mutants at 200mM [NaCl]. D) Structure of the LxCxE_WT_:Rb complex (PDB ID: 1GUX) depicting the changes in free energy of activation (*ΔΔG^‡^*). In panels A and C, LxCxE motif residues are represented on the x-axis, and mutations are indicated below each bar. Black bars: non-contact positions; Violet bars: contact positions; Filled bars: alanine scanning; Open bars: L22I, Y23F, C24S, Y25F, E26D, L28+3. Hatched bars: Y25H, L28+4. In panels B and D, residues are colored using a heat map depicting *ΔΔG^0,‡^* values in a scale from 0 (white) to 5 (red) kcal/mol.

In contrast, the activation barrier revealed a fairly flat energy landscape with negligible contributions near the N-terminus (D21-Y25) and moderate contributions (*ΔΔG**^‡^*** < 1 kcal/mol) near the C-terminus (Figure 2 C,D). We assessed the global features of the energy landscape using linear free energy relationship (LFER) plots, which quantify the proportion of the change in the free energy of binding explained by changes in the free energy of association^15,51,52^. The LFER plot for LxCxE_WT_ CFB was linear, which is a signature for the presence of one major rate-limiting barrier or TSE^15,51^ (SI Appendix, Figure S5). The slope of the LFER plot or α-value measures the position of the TSE along the reaction coordinate and takes values between 0 (early, substrate-like or disordered) and 1 (late, product-like or structured). The low α value (α= 0.15, SI Appendix, Figure S5) indicates that the TSE for LxCxE_WT_ CFB is disordered and located early in the reaction pathway, as reported for many CFB reactions (PDZ-IDR^23,53^, PTB-APP^27^, pKID-KIX^21^, TAZ1-IDR^22,24^ and PUMA-Mcl1^9^). Mutating L22 was more disruptive than mutating L28 at equilibrium, but only L28 had significant effects on the TSE, and these results were robust across different substitutions at both positions (Figure 2A-C). A similar asymmetry was present for charged residues D21 and E26 (Figure 2C). This suggests some degree of orientational constraints in the TSE, compatible with partial docking of the peptide to the L28 binding pocket, mirroring the results of other CFB reactions^32,53–55^. We conclude that LxCxE_WT_ CFB to Rb recapitulates multiple features of CFB reactions, making it a good model system to elucidate the energy landscape for CFB.

### Leffler analysis uncovers non-native energetics in LxCxE_WT_ CFB

Leffler plots correlate changes in activation (*ΔΔG*^‡^) and equilibrium (*ΔΔG*^0^) free energy of binding for each mutant, providing information on the energy landscape for CFB at single-residue resolution^52,56,57^. The energy landscape can in turn be interpreted in terms of the types of contacts formed, providing structural insight on the TSE. TSE energetics are often quantified using Φ-values (Φ = ΔΔG^‡^/ΔΔG^0^)^15,16^. While both Leffler plots and Φ-values report on TSE energetics, Φ-values become highly unreliable for small *ΔΔG^0^* values^18,56^ and a direct analysis using Leffler plots overcomes this limitation, enabling the identification of residues contributing to the TSE energetics even when *ΔΔG^0^* is small^56^.

Leffler plots can be divided into two areas corresponding to native-like or non-native energetics (Figure 3A and Materials and Methods). The native-like area of the Leffler plot (Figure 3A, white area) describes a smooth energy landscape where native contacts form gradually along the reaction coordinate. The non-native area of the Leffler plot (Figure 3A, light grey area) describes a rugged energy landscape characterized by the transient formation of contacts that are not present in the native state, also referred to as non-native contacts. To assess whether the energy landscape for CFB of LxCxE_WT_ to Rb was smooth or rugged we classified whether mutations had no energetic contribution (Figure 3A, closed black circles) or contributed native-like (Figure 3A, red points) or non-native (Figure 3A, green points) energetics in the TSE, using a conservative energy cutoff (0.2 kcal/mol) derived from the distribution of errors for *ΔΔG^0^* and *ΔΔG*^‡^ (see Materials Methods and SI Appendix Figure S6 and Table S4).

**Figure 3.**
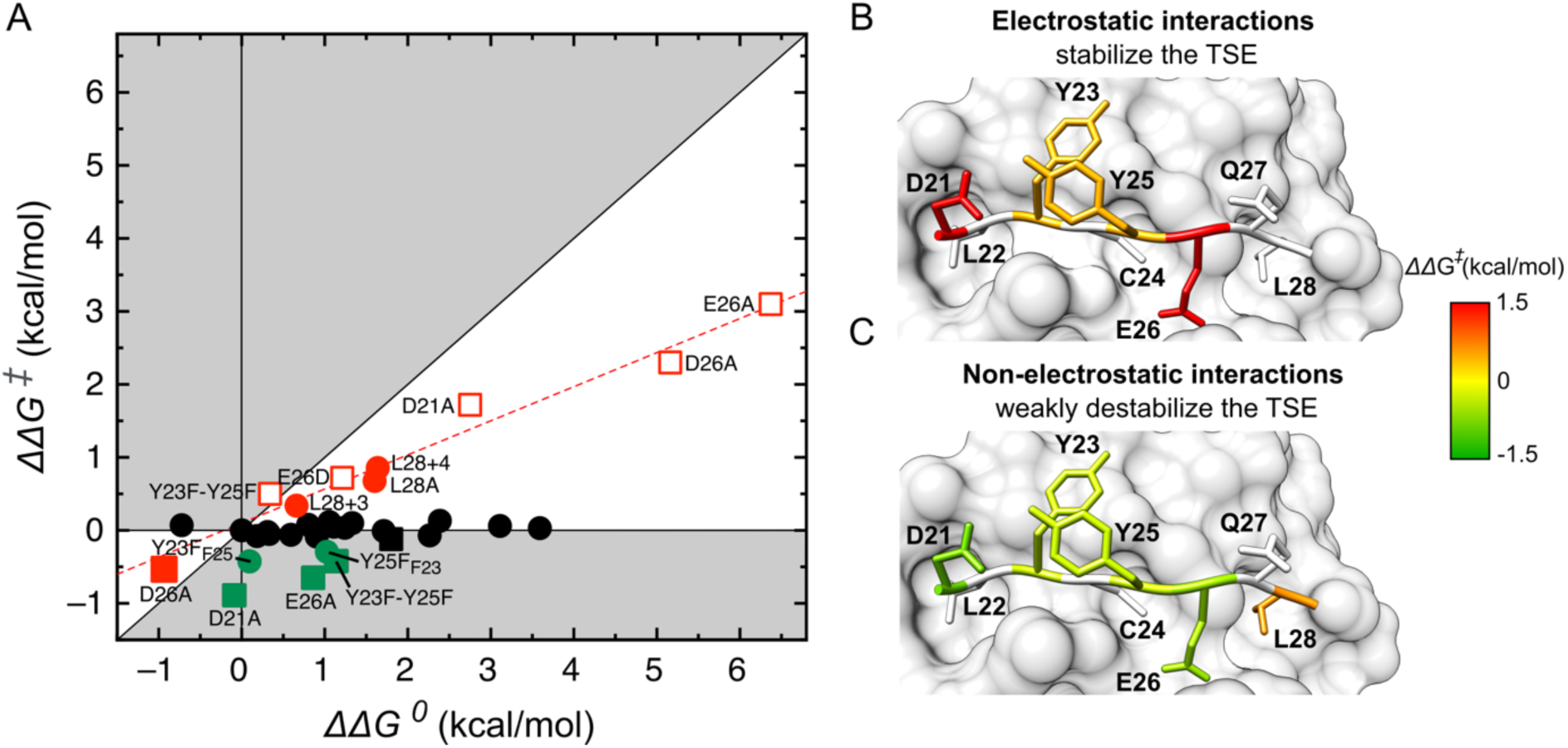
Leffler analysis of LxCxE_WT_ CFB using electrostatic dissection. (A) Leffler plot for the [Rb:LxCxE_WT_] complex depicting native-like (white) and non-native (grey) areas. Each point refers to a mutation classified as reflecting no energetic contribution (black circles), native-like energetics (red circles) or non-native energetics (green circles). Mutants for which electrostatic dissection was performed are depicted by squares, with electrostatic (empty squares, *ΔΔG^0^_E_* and *ΔΔG^‡^_E_*) and neutral (filled squares, *ΔΔG^0^_N_* and *ΔΔG^‡^_N_*) components shown. Black lines indicate the zero lines and the line with slope 1. Red dashed line: linear fit of mutations yielding native-like energetics (slope = 0.46 ± 0.03). (B-C) Residue-level decomposition of electrostatic (*ΔΔG^‡^_E_*, B) and non-electrostatic (*ΔΔG^‡^_N_*, C) energy contributions in the TSE. Y23 and Y25 values are those of the Y23F-Y25F mutant. For all other residues, values are those of the Ala mutation. *ΔΔG^‡^* values are represented as a heat map with stabilizing interactions depicted in red and destabilizing interactions depicted in green. The *ΔΔG^‡^_E_* values for D21A and E26A are beyond the upper limit of the scale (ΔΔG^‡^_E,D21A_= 1.72 kcal/mol and ΔΔG^‡^ _E,E26A_= 3.06 kcal/mol). For L28A the neutral component was obtained at 200 mM NaCl.

Alanine scanning identified native-like energetics compatible with a smooth energy landscape (Figure 2 C,D and SI Appendix, Table S2). However, Tyr-to-Phe mutations, which contribute to uncover non-native interactions^58^ by minimizing changes in the free energy of solvation (*ΔG_SOLV_*) or reorganization (*ΔG_REORG_*) ^27^ hinted at a more complex energy landscape. Several Tyr-to-Phe mutations decreased the overall stability but increased the association rate, indicating that they stabilize the TSE (SI Appendix, Table S2). The Leffler plot identified non-native energetics for Y23F-Y25F, Y23F_F25_ and Y25F_F23_ (Figure 3A, closed green circles). The latter two are derived mutants that assess the effect of the Y23F mutation on the F25 background (Y25F → Y23F-Y25F) and vice-versa. Many but not all of these effects were captured by Φ-value analysis (SI Appendix, Figure S7 and Table S2). The lack of correlation between *ΔΔG^‡^*and changes in hydrophobicity (R^2^ = 0.2) indicates that, as for other CFB reactions^27^, the energetics are not explained by an increase in nonspecific hydrophobic interactions due to the removal of polar moieties (SI Appendix, Supplementary Text).

### Electrostatics are responsible for compensatory energetics in the TSE for CFB

Electrostatic interactions between the negatively charged LxCxE_WT_ motif and the positively charged Rb LxCxE cleft speed up association through electrostatic steering (Figure 1C)^36^. The finding of non-native energetics prompted us to test whether other forces such as electrostatic interactions played additional roles in stabilizing the TSE for CFB of LxCxE_WT_ to Rb. To investigate this possibility, we assessed the dependence of the kinetic rate constants on ionic strength for LxCxE_WT_, Y23F-Y25F and the D21A, E26D and E26A charge mutants. All mutants showed two-state behavior at all ionic strengths, as reported for LxCxE_WT_ ^36^ (SI Appendix, Figure S8 and Tables S5-S10). We performed global fits for the dependence of the kinetic rate constants on ionic strength using the Debye-Hückel approximation^59,60^ (see Materials and Methods and SI Appendix, Figure S9) and calculated the electrostatic and neutral components for the change in the free energy of activation (*ΔΔG*^‡^*_E_* and *ΔΔG*^‡^*_N_*) and the free energy of binding at equilibrium (*ΔΔG*^0^*_E_* and *ΔΔG*^0^*_N_*) (SI Appendix Table S11). Using the E26A and E26D mutations, we also assessed the derived mutant D26A (E26D-E26A). This approach allowed us to map the neutral and electrostatic contributions of five mutations onto the Leffler plot (Figure 3A, empty and filled squares).

A striking pattern emerged upon performing the electrostatic dissection: for all mutants, the electrostatic component yielded native-like energetics (Figure 3A, open red squares). The stabilizing effect of electrostatic interactions from E26 was much larger (*ΔΔG^‡^* = 3.06 ± 0.11 kcal/mol) than detected without electrostatic dissection (*ΔΔG^‡^* = 1.01 ± 0.07 kcal/mol). The analysis identified new stabilizing interactions at positions D21, Y23 and Y25, and the Leffler plot reveals a conserved single slope of ∼0.4, indicating that all electrostatic interactions are developed to a similar degree in the TSE. Conversely, the assessment of neutral components confirmed the non-native energetics for positions Y23 and Y25 and uncovered additional non-native energetics at positions D21 and E26 (Figure 3A, closed green squares). All electrostatic components stabilize the TSE (Figure 3B), while most neutral components introduce energetic frustration in the TSE (Figure 3C). Therefore, electrostatic dissection appears as a powerful tool that localizes energetic frustration to neutral components and stabilizing energetics to electrostatic components.

### Ruling out ground state effects in CFB of LxCxE_WT_ to Rb

Ground state effects can confound Φ-value analysis and an accurate assessment TSE energetics if mutations change the free energy of the unbound state^45,46^, but they have been monitored for only a handful of CFB reactions (Supplementary Text)^9,21,55^. To rule out ground state effects, we assessed the conformation of Rb and the LxCxE motif mutants. The low RMSD value (0.53 Å) obtained when aligning the crystal structures of unliganded and LxCxE-bound Rb (SI Appendix, Figure S10) suggests that unbound Rb behaves as a fairly rigid receptor when binding to LxCxE_WT_. For LxCxE peptides, we first assessed secondary structure content using Far-UV circular dichroism (Far-UV CD) spectroscopy^61–63^. The Far-UV CD spectrum of most mutants was superimposable with LxCxE_WT_ and consistent with a highly disordered polypeptide with no persistent secondary structure, matching previous reports^43^ (SI Appendix, Supplementary Text and Figure S10). The changes in the CD spectrum for Tyr-to-Ala and Tyr-to-Phe mutants can be explained by the contributions from aromatic side chains to the Far-UV CD region^64,65^ (SI Appendix, Supplementary Text and Figure S10).

To rule out ground state effects for the Tyr-to-Phe mutants that revealed non-native energetics, we performed nuclear magnetic resonance (NMR) analysis. The ^1^H-^15^N HSQC spectrum of LxCxE_WT_ had low dispersion compatible with a disordered polypeptide (Figure 4A). The ^1^H-^15^N HSQC spectra of LxCxE_Y23F_, LxCxE_Y25F_ and LxCxE_Y23F-Y25F_ were superimposable with LxCxE_WT_ except for the N-terminal region (Q16-T20), where small chemical shift differences can be explained by the lack of N-acetylation on the LxCxE_WT_ peptide (Figure 4A). The ΔδCα - ΔδCβ secondary chemical shifts^66^ of LxCxE_WT_ displayed low values across all residues indicating a lack of secondary structure, a behavior that was mirrored by all Tyr-to-Phe mutants (Figure 4B). The elevated value for C24 likely reflects the influence of the two aromatic neighbors arising from known aromatic ring current effects on chemical shifts in disordered proteins^67^ and the Cβ value (∼27 ppm) corresponds to a reduced cysteine, ruling out cysteine oxidation. These results agreed with Cα secondary chemical shifts^68^ (SI appendix, Figure S11) and the helical content estimated from NMR and Far-UV CD experiments was low, ranging from 0.2 to 10 % depending on the method used (SI appendix, Table S12). Ground state effects could originate from disrupting pre-existing aromatic stacking present in the free LxCxE_WT_ peptide. However, the inspection of NOESY spectra of LxCxE_WT_ did not reveal any NOEs between aromatic ring protons of Y23 and Y25, indicating no significant population of Tyr stacking for unbound LxCxE_WT_ (SI appendix, Figure S11).

**Figure 4.**
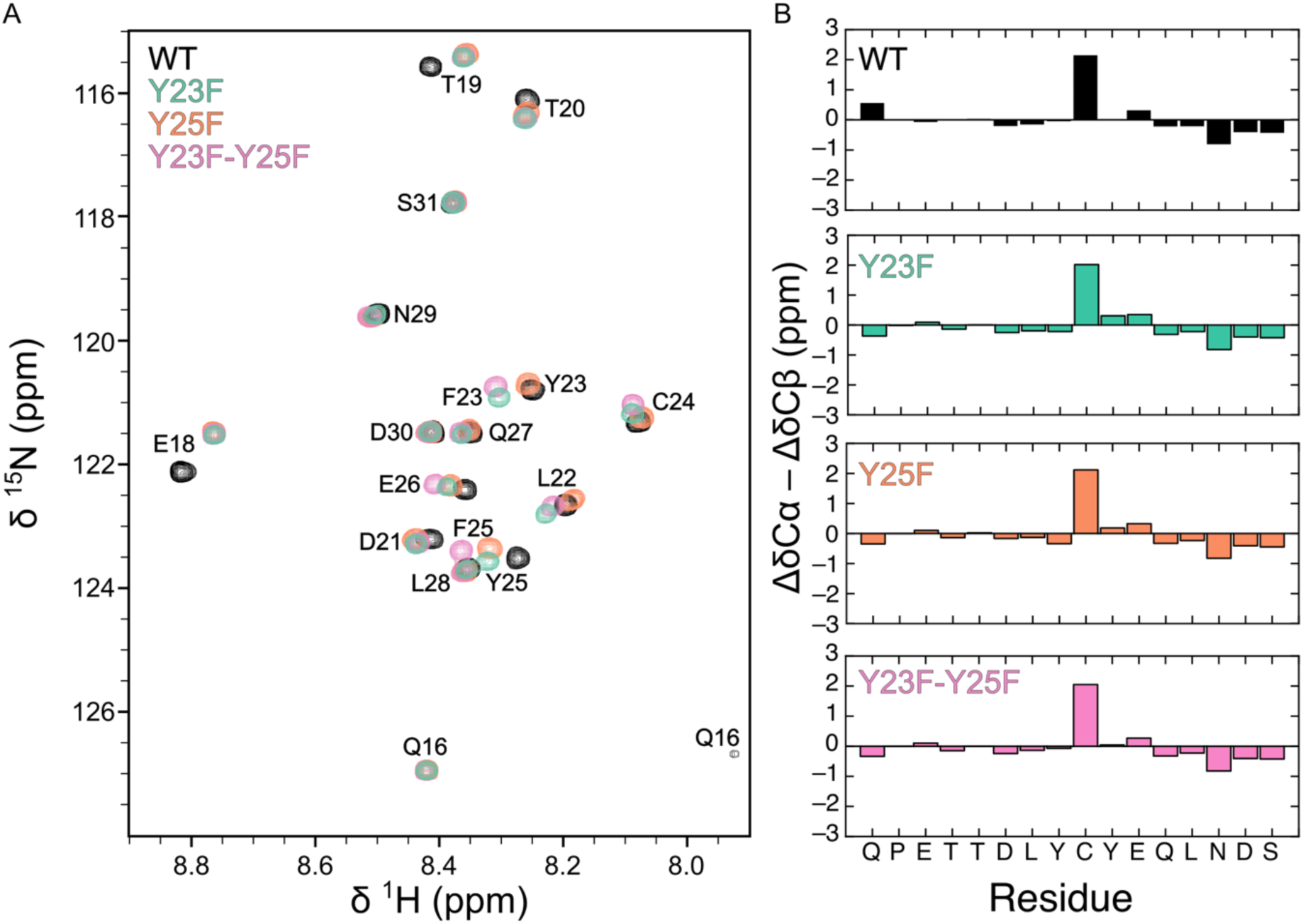
Tyr-to-Phe mutations do not change the conformational ensemble of the free LxCxE motif. A) ^1^H, ^15^N HSQC spectra of LxCxE_WT_ (black), LxCxE_Y23F_ (green), LxCxE_Y25F_ (orange) and LxCxE_Y23F-Y25F_ (pink). The spectra are mainly overlapping except the mutated residue/s. The differences between the spectra of LxCxE_WT_ and Y-to-F mutants at N-terminal residues (Q16, E18, T29 and S20) are explained by the fact that LxCxE_WT_ is not acetylated at its N-terminus. B) Secondary structure analysis of the LxCxE motif mutants using NMR. ΔδCα - ΔδCβ values for LxCxE_WT_ (black), LxCxE_Y23F_ (green), LxCxE_Y25F_ (orange) and LxCxE_Y23F-Y25F_ (pink). LxCxE_WT_ has negligible secondary structure content and the Y23F, Y25F and Y23F-Y25F mutations do not produce significant changes in secondary structure content.

Taken together, Far-UV CD and NMR experiments confirmed that LxCxE_WT_ and all mutants are highly disordered with no significant changes in secondary structure propensity or pre-existing intramolecular interactions, ruling out ground state effects for all mutants.

### Non-native energetics are a general feature of the TSE for CFB of IDRs

To assess whether non-native energetics were a conserved feature of CFB reactions, we performed a global analysis by selecting short IDRs that bound to folded domains through two-state reactions probed with extensive mutational scanning (see Materials and Methods). We compiled or computed *ΔΔG^0^* and *ΔΔG^‡^* values from literature for 177 mutations across five IDR complexes (S-peptide/S-protein, cMyb-KIX, pKID-KIX, APP-PTB^21,27,55,69^ and LxCxE_WT_- Rb) (Figure 5A). Only ∼20% of the mutations in this dataset were contributed by LxCxE_WT_-Rb (see SI Appendix, Supplementary Table S13). Mapping all mutations onto the Leffler plot revealed non-native energetics across all IDR complexes (Figure 5A and Figure 5B, black and open circles). More than half of the mutations revealing non-native energetics fail to be detected by Φ-value analysis using a standard cutoff (*ΔΔG^0^* > 0.6 kcal/mol) (Figure 5B, open circles), which explains in part why many were not detected previously. These results suggest that non-native energetics represent a common feature of the TSE for CFB of IDRs.

**Figure 5.**
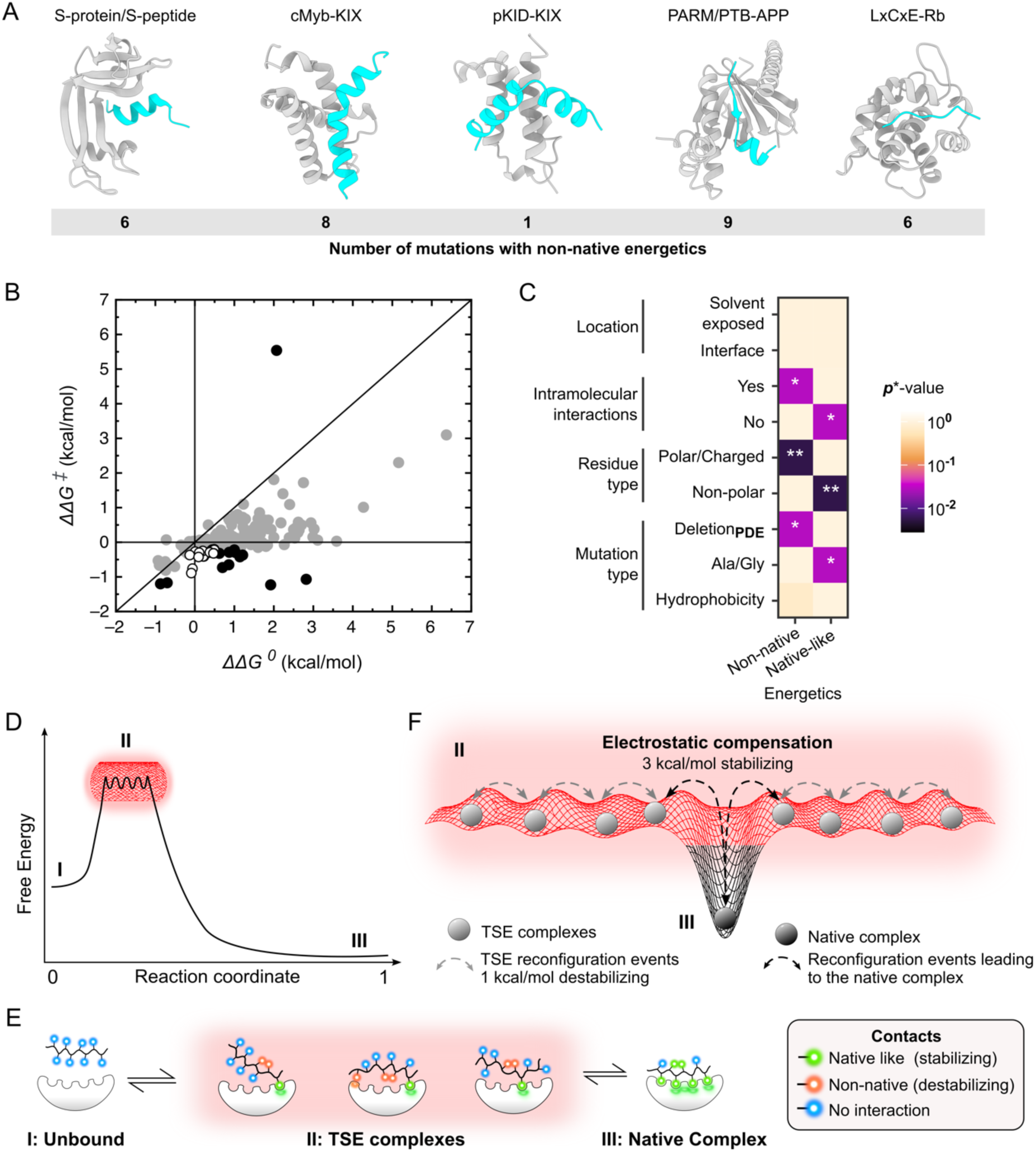
Global analysis of the energy landscape for CFB. A) Structure of the S-protein/S-peptide, cMyb-KIX, pKID-KIX, PARM/PTB-APP and LxCxE-Rb complexes^21,27,55,69^. B) Global Leffler plot (N = 177 mutations). Grey circles: Native-like energetics (N = 147); Black circles: Non-native energetics detected with Leffler and Φ-value analysis (N = 13); Open circles: Non-native energetics detected only with Leffler analysis (N = 17). C) Association between TSE energetics and residue-level features determined using a hypergeometric test corrected for the false discovery rate. Corrected *p\**-values are represented as a heat map, and statistical significance is indicated by asterisks: * = *p\** < 0.05; ** = *p\** < 0.01. D) Schematic representation of the energy landscape for CFB depicting the unbound (I), TSE (II) and native complex (III) states. The TSE is located early in the reaction coordinate and presents a rugged energy landscape. E) Microscopic view depicting the contact types detected in this work. The unbound IDR has no stable contacts. The TSE features multiple complexes in fast equilibrium (disordered) with partial docking of the IDR (native contact) and multiple non-native contacts. F) Close-up of the TSE (II) depicting a rugged and energetically frustrated energy landscape populated by small barriers created by non-native interactions. Stabilizing electrostatic interactions (∼3 kcal/mol) enable TSE reconfiguration events (∼1kcal/mol) to occur without complex dissociation. Some reconfigurations lead to the native complex (III). In panels D-F, electrostatic compensation is depicted as a red mesh.

Next, we asked whether non-native energetics were associated with specific residue-level features, including residue location, presence of intramolecular interactions, residue type and mutation class. Non-native energetics were associated with residues that establish intramolecular interactions (Figure 5C, *p\**<0.05) across most IDRs. The effect of disrupting hydrogen bond and stacking interactions in LxCxE_WT_ was recapitulated by disrupting helix-stabilizing salt bridges (R10, E2) or stacking interactions (F8, H12) in the S-peptide^55^ and H-bond interactions in the cMyb-KIX (T15 and N17)^69^ and pKID-KIX (R135)^21^ complexes (SI Appendix, Supplementary Table S13). This suggests that moieties involved in intramolecular interactions that fine-tune the binding affinity establish non-native contacts that create energetic frustration in the TSE for CFB.

Non-native energetics were also associated with the mutation of polar/charged residues (Figure 5C, *p\**<0.01) across all IDRs (SI Appendix, Supplementary Table S13). For cMyb-KIX (K3, S14, and N17)^69^ and PTB-APP (F764)^27^, where multiple mutations were performed at the same position, the mutation of polar/charged residues revealed either native-like or non-native energetics depending on the type of mutation performed. These results suggest that charged/polar residues establish non-native contacts, and may be responsible for compensatory energetics, across CFB TSEs. The Deletion_PDE_ class, which groups partial side chain deletions (P), double mutants (D) and electrostatic dissection (E), was associated with non-native energetics, while Ala-Gly scanning was associated with native-like energetics (Figure 5C, *p\**<0.05). This likely reflects the fact that while Ala-Gly scanning is a powerful approach to dissect contributions from binding versus secondary structure formation for helical IDRs^15,21,69^, full side chain deletions may obscure subtle energetics revealed when performing partial deletions^15,27^. These results underscore that Deletion_PDE_ mutations and sampling multiple mutations per position enable a comprehensive mapping of the TSE energetics^15,70^.

Taken together, the global analysis confirms and extends the findings of the LxCxE_WT_ model system, suggesting that non-native energetics represent a common feature of the TSE for CFB of IDRs. Non-native energetics are associated with intramolecular interactions that fine-tune the binding affinity and with charged/polar residues, which may be responsible for compensatory energetics in the TSE for CFB.

## DISCUSSION

In this study, we establish LxCxE_WT_ as a minimal model system for the CFB of IDRs. The rate limiting barrier for LxCxE_WT_ binding to Rb is located early in the reaction coordinate (α = 0.15) and indicates a highly disordered TSE (Figure 5D). This is consistent with all CFB reactions of IDRs that show no evidence of ground state effects^9,21–26^. The electrostatic dissection we performed revealed strong compensatory energetics that were otherwise obscured at 200mM NaCl (Figure 3). It localized stabilizing energetics to native-like electrostatic interactions and destabilizing energetics to non-native contacts of neutral nature. This indicates that a highly disordered TSE is coupled with a relatively rough energy landscape presenting non-native contacts that must reconfigure to allow the formation of the native complex (Figure 5D, E). Two hot spots for binding (L22 and C24) play no role in the TSE energetics, whereas positions that play a minor role in complex stability (e.g. D21) have strong contributions in the TSE (Figures 2 and 3). The fact that the TSE energetics are uncorrelated with the native state energetics provides additional evidence for the existence of reconfiguration events.

Electrostatic forces play key roles in stabilizing IDR complexes through electrostatic steering^10,11,48^ and tuning IDR affinities^71^, structural ensembles^72^ and interactions^73,74^. While protein folding, in general, is funneled by the native topology^20^, the TSE for CFB appears to present a relatively flat but rugged energy landscape featuring multiple shallow local minima that populate non-native configurations (Figure 5F). Electrostatics (Figure 5D-F, red mesh) could be envisioned to play a role akin to the reduction of dimensionality from a 3D to a 1D search in transcription factors binding to DNA^75^ by confining the configurational search space to the cognate binding pocket and preventing dissociation to allow non-native contacts to reconfigure. We term the mechanism through which electrostatic interactions offset energetic frustration in the TSE for CFB, electrostatic compensation. The partial docking/pre-orientation of the IDR with respect to the binding pocket (Figure 5E, this work and many IDRs^32,53–55)^ cooperates in reducing the dimensionality of the search for the native state. Non-native interactions are weakly destabilizing (*ΔΔG^‡^*∼1 kcal.mol^-1^) and are counterbalanced by stabilizing electrostatic interactions (*ΔΔG^‡^* ∼3 kcal.mol^-1^) (Figure 5F), suggesting rapid reconfiguration and a rapid search for native interactions. The exploration of non-native conformations in the search for the native state (Figure 5E,F) may explain the longer transition path times of IDR binding^29,30^ compared to protein folding^76^. This study provides the first robust experimental evidence for energetic frustration and reconfiguration events in the TSE for CFB, previously reported only in molecular dynamics studies of IDR binding trajectories^32^.

A global analysis of the energy landscape for CFB using Leffler plots (177 mutations)^21,27,55,69^ suggests that energetic frustration is a common feature of the TSE for CFB of IDRs (Figure 5 A,B). Non-native energetics were also detected upon mutating the folded domain, e.g. in the CFB TSE of PUMA-MCL-1^25^. Considering that partial deletions that minimize changes in reorganization and solvation free energy^27,77^ enabled the detection of non-native energetics for LxCxE_WT_ and were scarcely exploited in other systems, this global analysis likely represents a lower estimate for the prevalence of non-native energetics in this dataset. It further suggests that extensive mutational scanning^15,70^ and the dissection of electrostatic and neutral components will improve the detection of non-native and compensatory energetics in other IDRs. Initial analyses using ∼200 mutations^17,56^ captured global features of folding energy landscapes that were confirmed by larger datasets^18,19^. This underscores the potential our dataset to reveal general features of CFB energy landscapes.

IDRs can bind to different partners with high specificity and affinity. Many IDRs retain disorder when bound to their partners, and these fuzzy regions are highly frustrated^78,79^. This work identifies energetic frustration in the earliest steps of coupled folding and binding. It reveals that disorder and energetic frustration are features of the bottleneck of the reaction. The association of non-native energetics with interactions that increase complex stability (LxCxE_WT_^39,40,47^ and other IDRs, Figure 5C) suggests that energetic frustration may arise as a byproduct of optimizing the affinity of IDR complexes. Energetic frustration could also poise the system for maximal sensitivity, providing a mechanistic framework for understanding how subtle changes in the binding partner can template the folding of the IDR^14^. Electrostatic compensation is mediated by charged/polar residues, and many IDR complexes present complementary charges between the IDR and the binding cleft. This is an important experimental consideration since electrostatic compensation of non-native energetics may be present across the CFB of IDRs. This mechanism may enable fine-tuning of the affinity and specificity of IDR interactions, dictating functional selection for charge content within IDRs.

## ACKNOWLEDGEMENTS

L.B.C. and I.E.S. are Independent researchers from Consejo Nacional de Investigaciones Científicas y Técnicas (CONICET, Argentina). L.A. and J.G. were supported by a postdoctoral fellowship and N.A.G. by a doctoral fellowship from CONICET. The work was funded by Agencia Nacional de Promoción Científica y Tecnológica PICT No. 2019-02119 and No. 2021-01027 (to L.B.C.). This work was Funded by the European Union. Horizon Europe MSCA Staff Exchange project IDPfun - grant agreement No. 778247 (to L.B.C. and T.J.G.) Views and opinions expressed are however those of the author(s) only and do not necessarily reflect those of the European Union or the Research Executive Agency. Neither the European Union nor the granting authority can be held responsible for them. We are grateful to Mehrnoosh Arrar and Gary W. Daughdrill for critical reading of the manuscript. We thank to the Expression and Purification Core Facility at EMBL, especially Karine Lapouge, for support in CD measurements and Edward Lemke for providing access to Stopped Flow instrumentation.

## AUTHOR CONTRIBUTION STATEMENT

Conception/design: L.A., L.B.C.; Reagent generation: L.A., M.C. and P.S. Data acquisition: L.A., N.A.G, K.S., P.S.; Data analysis: L.A., K.S., J.G., L.B.C; Data interpretation: L.A., K.S., J.H, I.E.S., L.B.C.; Writing, original draft: L.A., L.B.C; Writing, editing, all authors; Funding acquisition: L.B.C. and T.J.G.; Supervision: D.W., J.L., J.H., L.B.C.

## DATA AVAILABILITY

All the raw data underlying Main Figures and Extended Data is available as Source Data Files. Raw NMR data are available from the authors upon request.

## MATERIAL AND METHODS

### Protein Expression and Purification

The human retinoblastoma protein (Uniprot ID: P06400) AB domain (372–787 aa) with a deletion of the loop connecting the A and B pockets (aa. 372-581 and 643-787), here named Rb, was expressed and purified as previously described^36,38,41^. Briefly, Rb was recombinantly expressed from a pRSET-A vector containing an N-terminal 6xHis-tag. *Escherichia coli* C41 cells transformed with this vector were grown to an OD_600_ of 0.6-0.8 and induced with 1 mM isopropyl β-D-1-thiogalactopyranoside (IPTG), with the protein expression taking place overnight at 18°C. Cells were pelleted and resuspended in a lysis buffer containing 50 mM Tris:Cl, 1 mM EDTA, 2 mM β-mercaptoethanol and 2 mM PMSF (pH 7.5). Sonication was performed to lyse the cells and following centrifugation, the resulting soluble fraction was purified using a Ni^2+^ -nitrilotriacetic acid immobilized metal affinity chromatography (IMAC) followed by a sulphate cation exchange (SP-Sepharose) chromatography. To remove the His tag, the protein was cleaved by treatment with Thrombin (Sigma-Aldrich, St. Louis, MO, USA) protease overnight at 4°C. The protein was subjected to size-exclusion chromatography (Superdex 75) to remove the cleaved His tag and achieve the final pure sample. Rb was quantified by absorbance at 280 nm using a molar extinction coefficient of 40,340 M^-1^ cm^-1^ and stored in 20 mM sodium phosphate buffer, 200 mM NaCl and 2 mM dithiothreitol (DTT), 10% glycerol at -80°C.

### Peptide synthesis and labeling

Peptides corresponding to residues 16-31 from HPV16 E7 LxCxE (Uniprot: P03129) were synthesized by the chemical biology core facility of EMBL, Heidelberg, Germany. The assembly of all linear protected peptides was performed automatically by solid-phase peptide synthesis (SPPS) using the standard 9-fluorenylmethoxycarbonyl/tertiobutyl (Fmoc/tBu) protection strategy. Automated synthesis was performed with a Biotage Syro I peptide synthesizer. Relative to the resin loading, coupling reactions were performed using: 5 eq. of Nα-Fmoc-protected amino acid (0.4mol.l-1), 5 eq. PyBOP (0.4 mol.l-1) dissolved in DMF and 10 eq. DIPEA (1.95 mol.l-1) dissolved in NMP for 40 min. The resin was then washed three times with DMF (10 mL/g resin) for 1 min. Nα-Fmoc protecting groups were removed by treatment with piperidine/DMF (4:6) (20mL/g resin) for 3 min. The process was repeated three times and the resin was further washed six times with DMF (15 mL/g resin) for 1 min. After the synthesis, all the compounds were acetylated on the N-term using a solution of acetic anhydride/Et3N/DCM (1/2/7) for 10 min twice. For FITC compounds, 1.5 eq FITC was added to 1eq of free N-term peptide (20-30mg) dissolved in 250 uL DMF containing 5 eq. DIPEA (pH>10); Reaction time 30 min. Deprotection of peptides from the resin was done using a solution of TFA/TIS/H2O (95/2.5/2.5) for 2h. Peptides were finally purified on reverse phase HPLC and analyzed on an LC Agilent 1290 series equipment and High-resolution mass spectra (HR/MS) were obtained at the metabolomics core facility of EMBL Heidelberg. Peptides sequences can be found in SI Appendix, Table S1.

The WT, L22I, Y25H, and L28+4 peptides were purchased from Genscript (New Jersey, USA) with purity greater than 95%. In those cases, FITC labeling at the free N-terminus was carried out in 100 mM sodium carbonate buffer, pH 8.0, at room temperature overnight, and labeled peptides were separated from free FITC by a PD-10 column (GE Healthcare, Uppsala, Sweden). The buffer used for elution was 50 mM Tris:HCl, pH 7.5. The concentration of all preparations was judged as stated above using the reported extinction coefficients for tyrosine and tryptophan in buffer solution or by UV absorbance at 220 nm in HCl. The FITC concentration was determined at pH 7.0 and 495 nm using a molar extinction coefficient of 75,000 M^-1^ cm^-1^.

### Chemicals and solutions

All chemical reagents were purchased from Sigma-Aldrich, and all solutions were prepared with distilled and deionized (Milli-Q) water and filtered through 0.22 μM membranes before use. All measurements were performed in a standard buffer containing 20 mM sodium phosphate buffer (pH 7.0), 200 mM NaCl and 1 mM DTT at 20.0°C, unless stated otherwise.

### AlphaScreen assay

To obtain equilibrium dissociation constants, an AlphaScreen™ assay was used as previously described^49^. In brief, the inhibition of the interaction between a biotinylated LxCxE-peptide (Biotin-QPETTDLYSYEQLNDS) and Rb with N-terminal 6xHis-tag was measured with Streptavidin-coated Donor beads and Nickel Chelate-coated Acceptor beads. The AlphaScreen signal was measured in an Envision plate reader (Perkin Elmer, United States of America) and IC_50_-values were calculated in GraphPadPrism using the model/equation log(Inhibitor) vs. response with variable slope. The equilibrium dissociation constants were also obtained from the AlphaScreen assay by fitting the data considering the binding at equilibrium of both the Biotin-LxCxE peptide and the inhibiting peptides according to a previously described model^80^.

### Kinetic measurements

The kinetic experiments were performed using stopped-flow fluorescence spectroscopy (SFM-3000, Bio-Logic, France). The excitation monochromator was set to 495 nm with a band-pass 8 nm, and emission was collected through a 515 nm cut-off filter (ThorLabs, New Jersey, United States). All concentrations reported are those of the measurement cell, which resulted from mixing different volumes of protein, peptide and buffer solutions from each syringe. The association and dissociation reactions were monitored by following the fluorescence intensity change of the FITC moiety coupled to the LxCxE peptides at 20.0 °C.

### Pseudo-first-order association kinetics

For the association reactions, a series of eight measurement points were performed where FITC-peptide concentration was held constant and Rb concentration was sequentially increased. Pseudo-first order conditions were used with respect to Rb, using a starting concentration of Rb at least five times higher than the concentration of the FITC-peptide and high enough to ensure a good signal-to-noise ratio (SI Appendix, Table S3). For each mutant, the measurement time base was chosen to record data for at least 5 times the half-life time of each decay. For each measurement point, 6 to 11 traces containing at least 20000 measurements were averaged. The traces were fit to a single exponential function (Eq. (1)) to obtain the observed rate constant (*k_obs_*):

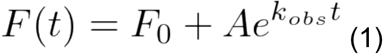

where A is the amplitude, F_0_ is the initial fluorescence value and *k_obs_* the observed rate constant. In all cases the data could be fitted to a single exponential function. The high affinity and good signal-to-noise ratio of the fluorescence signal enabled the acquisition of high-quality kinetic data for all but one of the mutants (Y23A-Y25A).

Under pseudo-first order conditions, the following relationship holds:

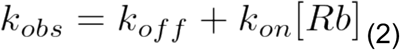

Therefore, the association rate constant, *k_on_*, was obtained from the slope of the linear fit of the plot of *k_obs_* as a function of Rb concentration using Eq. (2). In order to avoid underestimating the true error in the experiment, a 5% uncertainty in the determination of the Rb concentration was included in the error analysis for the calculation of *k_on_*.

### Dissociation kinetics

In principle, the *k_off_* value can be obtained from Eq. (2), but this determination, being an extrapolation to the x-axis intercept, is prone to error when the value of *k_off_* is small. For this reason, the dissociation traces were independently estimated by displacing a stoichiometric complex of Rb and FITC-labelled peptide with a 20-fold excess of unlabelled peptide at concentrations where a fraction of complex equal or higher to 0.34 was obtained to ensure a high enough signal to noise ratio (SI Appendix, Table S3). The estimated dissociation rate constants (*k_off_*) were obtained by fitting the observed fluorescence traces to a mono-exponential decay (Eq. (1)).

It is important to note that in our case, the *k_on_* and *k_off_* values were obtained from two independent orthogonal experiments. Thus, there is no correlation term between the estimates of *k_on_* and *k_off_*^81^. The excellent signal to noise ratio in our system allowed us to determine *k_on_* and *k_off_* values with high precision when fitting data from kinetic traces and PFO plots. Even when accounting for a 5% error in the determination of Rb protein concentration, this translated into highly precise estimation of *ΔΔG^‡^*, *ΔΔG^0^* and Φ-values, with all errors being smaller than 0.2 (SI Appendix, Figure S6) even for *ΔΔG^0^* < 0.6 kcal/mol (see section for Φ-value analysis).

### Energetic changes of point mutations

To assess the effects of mutations on the energetics of binding, we defined the changes in equilibrium (*ΔΔG^0^*) and activation (*ΔΔG^‡^*) free energy of CFB for each mutant with respect to LxCxE_WT_.

Specifically, *ΔΔG^0^* is the free energy change between products and reactants of the mutant minus that of LxCxE_WT_:

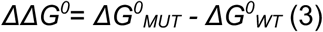

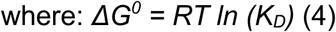

Similarly, *ΔΔG^‡^* is the free energy change between TSE and reactants of the mutant minus that of LxCxE_WT_ :

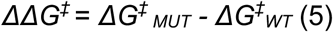

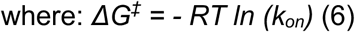

### NaCl dependence of the equilibrium and kinetic rate constants

The dependence of the equilibrium and kinetic constants on NaCl concentration was assessed by performing association and dissociation experiments as described above, using reaction buffer [20 mM sodium phosphate (pH 7.0), 1 mM DTT] supplemented with varying concentrations of NaCl ranging between 0.08 and 0.68 M (SI Appendix, Table S5). At each salt concentration, the time base was chosen to record data for at least 5 times the half-life time of each decay. Kinetic dissociation traces were recorded at each [NaCl] and *k_off_* values were calculated directly from fitting the fluorescence dissociation traces to a mono-exponential function as described above. The association rate constant *k_on_* at each [NaCl] was calculated from one association experiment as *k_on_*=(*k_obs_-k_off_*)/[Rb] using Eq. (1) and the *k_off_* values determined independently at each NaCl concentration considering a 5% error in the determination of Rb concentration. Kinetic traces were mono-exponential at all NaCl concentrations tested (SI Appendix, Figure S8). Standard deviations for each kinetic constant were calculated by error propagation of the standard deviations of fitted parameters. All values are reported in SI Appendix, Table S6-10.

The magnitude of variation in the equilibrium and rate constant produced by variations in [NaCl] can be characterized by the slope of –log *K_D_*, log *k_on_* and log *k_off_* versus log [NaCl], also referred as Γ_EQ_, Γ*_on_* and Γ*_off_*.

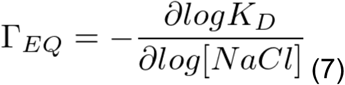

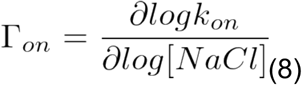

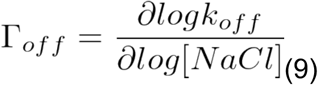

All values are reported in SI Appendix, Figure S7.

### Analysis of the contribution of neutral and electrostatic interactions in the TSE for binding using charge mutants

The neutral and electrostatic components of the change in free energy of binding for mutations of charged or polar residues were dissected using Debye-Hückel approximation which describes the dependence of the association rate constant *k_on_* with ionic strength. This model assumes that *k_on_* is correlated with the electrostatic interaction energy between the proteins, which is altered by the ionic strength of the solution. This dependence has been determined empirically by Vijayakumar et *al*.^59^ to follow the relationship:

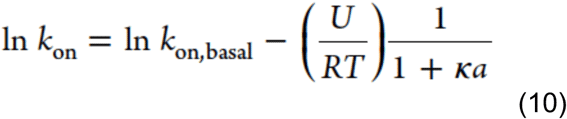

where *k_on,basal_* (units: M^-1^s^-1^) represents the basal *k_on_* value in the absence of long-range electrostatic interactions (in the experimental scenario, at infinite ionic strength, i.e. at maximal electrostatic screening by salt). The variable *U* (units: kcal/mol) is the electrostatic interaction energy between the associating proteins^82^ and the variable *a* is the minimal distance of approach between oppositely charged residues^10^. *k_on,basal_*, *U*, and *a* are fitting parameters, *R* is the gas constant, and T is the temperature. *κ* is the Debye-Hückel screening parameter which relates to the ionic strength of the solution. It is given by:

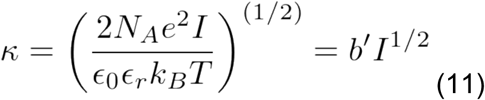

where *N_A_* and *k_B_* are the Avogadro and Boltzmann constants, respectively, ε_0_ and ε_r_ are the electrical permittivity of the vacuum and the dielectric constant of water, and *I* is ionic strength. Therefore, Eq. (6) can be rewritten as a function of ionic strength as:

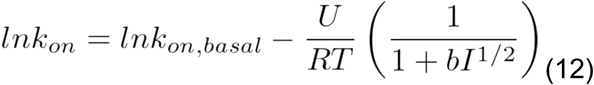

where *b* = *b’\**a. The parameter *a* was initially defined as the distance between oppositely charged residues on the surface of both proteins in the transition state^10^ and later reinterpreted as the sum of the radii for the two associating proteins^59^. Schreiber and Fersht, in their study of the proteins barnase and barstar^10^, found a strong dependence of *k_on_* with I, with the best fit for their system obtained using a value of 5.6 Å for the *a* constant (see Eq. (12)). Subsequently multiple studies applied the same approach, by fixing the *a* constant to a value of 6 Å (^48,82–84)^. This formalism can also be used to model the dependence of the dissociation rate constant, *k_off_*, with ionic strength (^85^).

To apply this model, we calculated the *ln k_on_* and *ln k_off_* values for each mutant at each ionic strength from the measured *k_on_* and *k_off_* values (SI Appendix, Figures S7-S8 and Tables S6-S10). We then globally fitted the dependence of the association and dissociation rate constants with ionic strength for all mutants using Eq. (12) and the values of ln *k_on, basal_* and U*_on_*/RT or those of ln *k_off, basal_* and U*_off_* /RT were obtained as fitting parameters for each mutant. The constant *b* was fixed considering a value of the parameter *a* of 6 Å (SI Appendix, Table S11).

### Analysis of the Transition State Ensemble: Leffler plots and Φ-value analysis

#### Leffler analysis

The structure of the transition state ensemble (TSE) was probed using Leffler plots^52^, which correlate the changes in activation (*ΔΔG^‡^*) and equilibrium (*ΔΔG^0^*) free energy of binding for each mutant. Leffler plots can be divided into two areas corresponding to native-like or non-native energetics. The native-like area of the Leffler plot (white area) describes a smooth energy landscape defined by the gradual formation of native contacts along the reaction coordinate, evidenced by mutations whose effect on the TSE is of equal or lower magnitude than at equilibrium (|*ΔΔG^‡^*| *≤* |*ΔΔG^0^*|). Within this area, *ΔΔG*^‡^ = 0 (Φ = 0) identifies residues that do not form interactions in the TSE, *ΔΔG^‡^* = *ΔΔG^0^* (Φ = 1) identifies residues that form native contacts in the TSE and intermediate values (|*ΔΔG^‡^*| *<* |*ΔΔG^0^*|, 0< Φ < 1) identify residues with a fractional population of native contacts in the TSE. By contrast, the non-native area of the Leffler plot (light grey area) describes a rugged energy landscape defined by mutations whose effect on the TSE is of opposite sign to (*ΔΔG^‡^* > 0 & *ΔΔG^0^* < 0 or *ΔΔG^‡^* < 0 & *ΔΔG^0^* > 0, Φ < 0) or by mutations whose effect exceeds that at equilibrium (|*ΔΔG^‡^*| *>ΔΔG^0^*|; Φ > 1). Non-native energetics are commonly interpreted to reflect the formation of contacts that are not present in the native state, also called non-native contacts.

Based on the distribution of errors for *ΔΔG^0^*and *ΔΔG*^‡^, which showed average standard deviation values < 0.1 kcal/mol (SI Appendix, Figure S6), we chose |*ΔΔG*^‡^| > 0.2 kcal/mol to classify whether a given residue had no energetic contribution, contributed stabilizing native-like energetics or non-native energetics in the TSE (SI Appendix Table S4). For non-native energetics where *ΔΔG^‡^* and *ΔΔG^0^* were of the same sign, we required that, |*ΔΔG^‡^*| exceeded |*ΔΔG^0^*| by more than 0.2 kcal/mol. The classification of all mutations using Leffler analysis is provided in SI Appendix, Table S4.

#### Φ-value analysis

The effect of a given mutation on the TSE for binding can be quantified by its Φ-value, which is calculated as the ratio between *ΔΔG**^‡^*** and *ΔΔG^0^*:

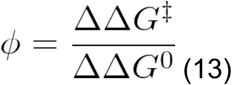

In the simplest case, the Φ-value can take values between zero and one, and represents an index of native-like structure of the mutated residue in the transition state^15,16,56^. Moreover, Φ-values between zero and one are indicative of a smooth energy landscape for binding with gradual development of native interactions along the reaction coordinate (white area of the Leffler plot) and Φ-values that are negative or larger than one reveal non-native energetics which may indicate the existence of non-native interactions that precede the formation of native contacts (grey area of the Leffler plot)^15,16^. Φ-value analysis is a useful tool for mapping TSE for folding and binding reactions, but because the Φ-value is defined as the ratio between *ΔΔG**^‡^*** and *ΔΔG^0^*, Φ-values calculated for mutations producing small changes in overall stability (*ΔΔG^0^*) are subject to large errors, and there has been debate about the minimum *ΔΔG^0^* value that warrants an accurate estimation of Φ-values^15,56^. Sanchez et *al.* proposed that Φ-values are accurate when *ΔΔG^0^* ≥ 1.67 kcal/mol^56^ while a less stringent criterion by Fersht et *al.* considered Φ-values reliable when *ΔΔG^0^* ≥ 0.6 kcal/mol^15^. The determination of Φ-values with higher precision in folding and binding reactions (SD < 0.2 kcal /mol) increases their reliability, which led some authors to report Φ-values for mutations with *ΔΔG^0^* < 0.6 kcal /mol^25,26^.

In this work we inform Φ-values when *ΔΔG^0^* ≥ 0.6 kcal /mol. Errors in the Φ-values were calculated using standard error propagation methods directly from Eq. (13) as previously performed by Sato et *al.*^86^. The errors for all Φ-values were < 0.2 (SI Appendix, Figure S6). The classification of all mutations using Φ-value analysis is provided in SI Appendix, Table S4.

#### Calculation of Φ_n_ and Φ_e_ values

The Φ*_N_* value was calculated following Eq. (13) as:

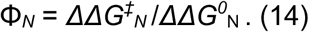

*ΔΔG**^‡^ _N_*** was calculated as:

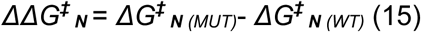

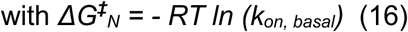

using the *k_on, basal_* value obtained from fitting LxCxE_WT_ and the mutants to the Debye-Hückel model.

*ΔΔG^0^ **_N_*** was calculated as:

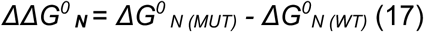

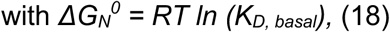

where *K_D, basal_ = k_off, basal_ /k_on, basal_*.

Since *U* describes the electrostatic interaction energy between the associating proteins^85^, the values of U_on_ for LxCxE_WT_ and each mutant obtained from the fitting of Eq. (8) were used to calculate *ΔΔG**^‡^*** _E,_ as *ΔΔG**^‡^*** _E_ = U_on *(MUT)*_ - U_on *(WT)*_.

The *ΔΔG^0^_E_* for LxCxE_WT_ and each mutant was calculated using U_on_ and U_off_ (also obtained as a fitting parameter) as *ΔΔG^0^_E_* = U_off_ - U_on._ Then, Φ*_E_* was calculated following Eq. (13) as Φ*_E_* = *ΔΔG**^‡^*** _E_ /*ΔΔG^0^* _E_. SI Appendix, Tables S11 summarize the results for each mutant.

### Far-UV Circular Dichroism (CD) spectroscopy

#### Measurement of Far-UV CD spectra

CD measurements were carried out on a J-815 CD spectropolarimeter equipped with a Peltier thermoregulation system (Jasco, Easton, PA, USA) set at 20°C. CD spectra were recorded between 190 and 260 nm at standard sensitivity and 50 nm/min scanning speed with a data pitch of 0.1 mm and a bandwidth of 1 mm using a quartz cuvette of 0.2 mm. All spectra were an accumulation of 10 scans. Samples were diluted to concentrations in the range of 0.4–0.6 mg/mL in 20 mM sodium phosphate, pH 7.0, and DTT 1 mM.

Mean residue weight (MRW) was calculated by the molecular weight divided by the number of amino acids. The mean residue ellipticity (MRE) was calculated from the Eq. (19):

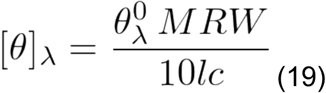

where *l* is the path length of the cuvette, and *c* is the concentration given in grams/ml.

#### Calculation of helical content from CD measurements

The ellipticity at 222 nm is assumed to be linearly related to the average helix content (*f*_H_). This can be calculated using two different methods. The first method^61^ uses the following empirical relationship:

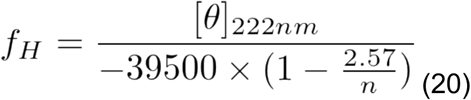

which calculates the ratio between the experimental molar ellipticity value at 222 nm and the molar ellipticity value corresponding to 100% helical content, where n is the number of residues in the peptide.

The second method^87^ uses the baseline ellipticities of both the random coil (θ_C_) and the complete helix (θ_H_):

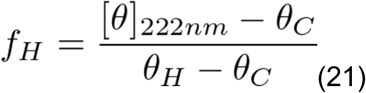

The values of θ_C_ and θ_H_ depend on the temperature and are given by the following expression:

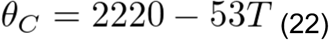

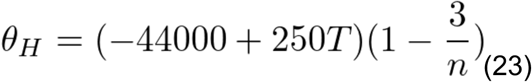

where T is the temperature in °C and n is the chain length in residues. The percentage of helix was calculated as f_H_ * 100.

### NMR spectroscopy

NMR experiments for the LxCxE_WT_, LxCxE_Y23F_, LxCxE_Y25F_ and LxCxE_Y23F-Y25F_ peptides were performed at a calibrated temperature of 283 K on a 600 MHz Bruker Avance II+ spectrometer with a room temperature probe (homonuclear ^1^H, ^1^H COSY, TOCSY, and NOESY experiments) and a 700 MHz Avance IIIHD spectrometer with a TCI cryoprobe (^1^H, ^13^C, and ^1^H, ^15^N HSQC). NMR samples contained 0.7-1.3 mM peptide in sodium phosphate 20 mM, pH 6.0, 75 mM NaCl, 5 mM TCEP and 10% (v/v) D_2_O. Assignments were done from ^1^H-^15^N-HSQCs, ^1^H-^15^N-COSY and ^1^H-^15^N-TOCSY. NMR spectra were analyzed using the NMRViewJ^88^ software. All residues were assigned except proline 17. Random coil chemical shifts were predicted using each peptide sequence and the web application of the University of Copenhagen (https://www1.bio.ku.dk/english/research/bms/sbinlab/randomchemicalshifts1/) according to Kjaergaard et al^89,90^. The secondary chemical shifts and per-residue helical content of each peptide were further analyzed using the δ2Δ^91^ and CSI 3.0 tools^92^.

### Comparison with previous studies

We conducted a global analysis of the TSE for CFB of other IDRs to folded domains to assess whether the features observed for the LxCxE motif were common to the TSE for CFB of other IDRs. We limited our analysis to systems in which the IDR was shorter than 35 residues and for which extensive mutagenesis was performed, followed by kinetic analysis using stopped-flow spectrometry. Based on these criteria, we selected the following systems: S-peptide-S-protein^55^, pKID-KIX^21^, cMyb-KIX^69^, and PTB-APP^27^ in addition to the LxCxE motif:Rb complex (this work). For each system, we calculated *ΔΔG^0^* and *ΔΔG**^‡^*** values from the published kinetic data for all mutations. Using this dataset, we classified each mutation according to a TSE energetics category and four residue-level categories as explained below (SI Appendix, Table S13). For the TSE energetics we performed a binary classification of the energetics as “non-native” or “native-like” depending on whether they fell in the “non-native” or in the “native-like” area of the Leffler plot (corresponding to Φ-values between zero and one). Second, we used the available PDB structures (PDB ID: 2RLN, 1KDX, 1SB0, and 3SUZ for S-peptide-S-protein, pKID-KIX, c-Myb-KIX and PTB-APP, respectively) to classify all mutated residues according to the following residue-level features: Residue location: Interface (side chain establishes contacts with the globular partner) or Solvent (side chain is exposed to solvent and does not establish contacts with the globular partner); Intramolecular interactions: classifies whether the mutated residue forms intramolecular interactions in the bound state; Residue type: residues were classified into polar/charged or non-polar; Mutation type: We classified the type of mutation into Ala/Gly (replacement by alanine or glycine), Deletion_PDE_ (partial side chain deletion, double mutant or electrostatic dissection) or Hydrophobicity (replacement by a side chain that increases hydrophobicity). We only considered standard mutations, excluding post-translational modifications.

#### Statistical analysis of association between TSE energetics and features of mutations

We assessed potential associations between (A) the TSE energetics category (“Non-native” or “Others”) and (B) four types of residue-level features, which included: B1: Location of the mutated residue (“Solvent exposed” or “Interface”); B2: presence of intramolecular interactions (“Yes” or “No”); B3: Type of mutated residue (“Polar/charged” or “Non-polar”) and B4: Type of mutation (“Deletion_PDE_”, “Deletion”, and “Hydrophobicity”).

Associations were assessed using a hypergeometric test as performed previously^93^. Briefly, we considered N mutations, for which the Energetics category A was observed in *a* mutations, the feature category B was observed in *b* mutations, and both categories A and B were observed together in *z* mutations, with z ≤ min (a,b). The hypergeometric test quantifies the probability of observing *z* or more successes (co-occurrences) when sampling without replacement *b* elements from a population of size N containing *a* successes. The *p*-value is defined as the sum of the probabilities of observing z or more successes. We applied the Benjamini-Hochberg^94^ correction for multiple comparisons to control for the false discovery rate. A positive association is reported if the corrected *p*\* value is smaller than the chosen cutoff (*p\** < 0.05). The number of mutations considered was N=177 and the number of comparisons was 18 (Energetics x B1 = 4 comparisons; Energetics x B2 = 4 comparisons; Energetics x B3 = 4 comparisons and Energetics x B4 = 6 comparisons). All calculations were performed using the phyper function available in the stats R library^95^.

### Change in hydrophobicity

The partition coefficient (P) measures the degree to which a compound dissolves in water versus an organic solvent. In particular, P is calculated as the ratio between the concentration of the molecule dissolved in the organic phase and the concentration of the molecule dissolved in the aqueous phase. Although P is a constant, its value is dependent on the choice of the organic solvent and on the conditions of measurement. ChemDraw from PerkinElmer Informatics was used to obtain the values of P for the different residues. This program calculates the Ghose-Crippen octanol-water partition coefficient of the desired compound (in this case each amino acid)^96^. For each mutation, the change in hydrophobicity was expressed as Δlog P, calculated as Δlog P = log P_MUT_ - log P_WT_, following ^27^. For more information regarding this topic see SI Appendix, Figure S12.

### Data analysis and fitting

Fitting was carried out using the Profit software (Quantumsoft, Zurich, Switzerland) to obtain parameters and their standard deviations. Protein complex structures were visualized and rendered using UCSF Chimera^97^ and PyMOL^98^ for molecular visualization and figure preparation.

## Notes

### Competing Interest Statement

The authors have declared no competing interest.

### Summary of Updates

We have expanded the global analysis of the energy landscape for coupled folding and binding and include a working model for the proposed mechanism.

