## Supplementary Material for "Electrostatic interactions compensate energetic frustration in the coupled folding and binding of intrinsically disordered proteins"

### Supplementary Information

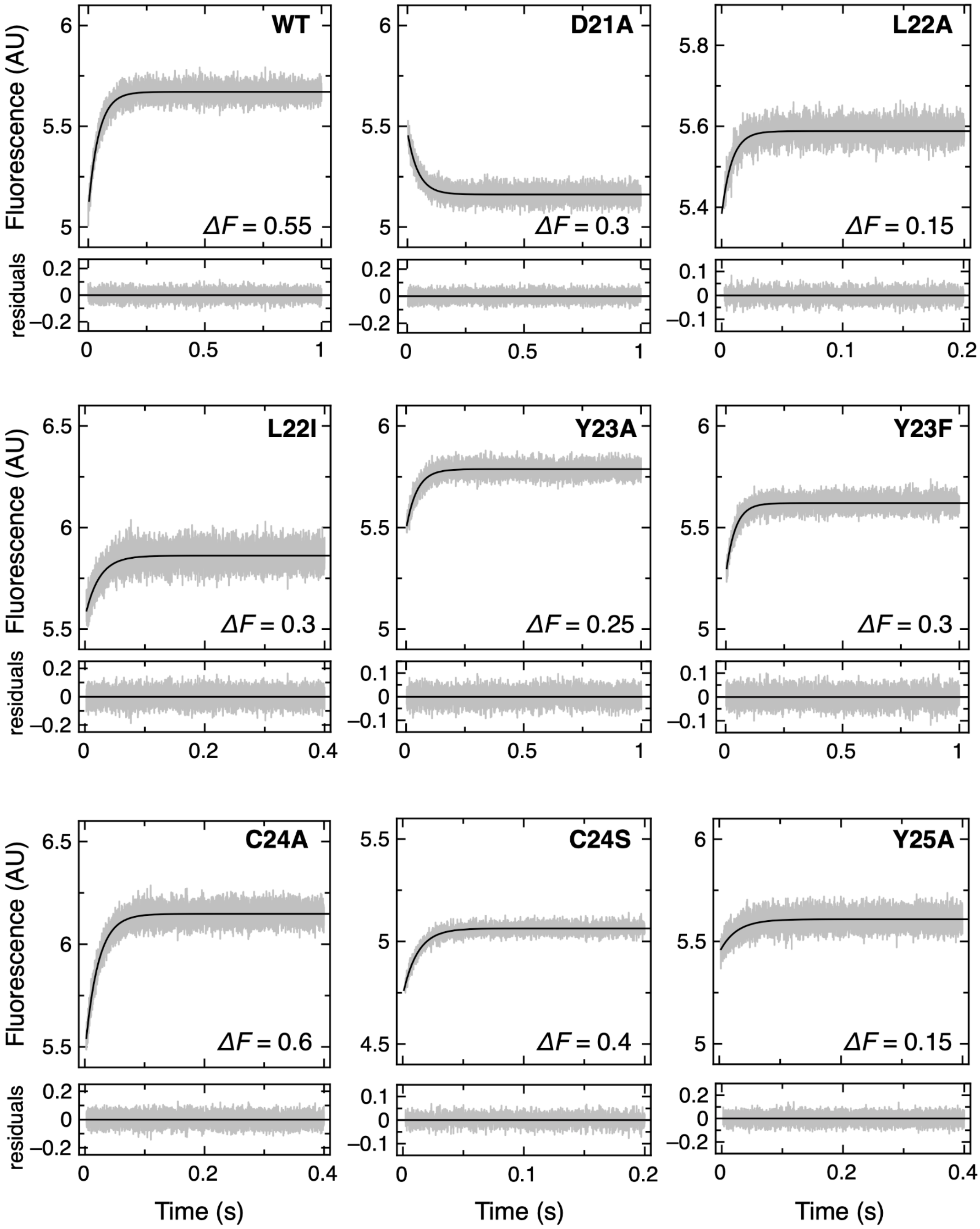
WT
ΔF = 0.55
D21A
ΔF = 0.3
L22A
ΔF = 0.15
L22I
ΔF = 0.3
Y23A
ΔF = 0.25
Y23F
ΔF = 0.3
C24A
ΔF = 0.6
C24S
ΔF = 0.4
Y25A
ΔF = 0.15
Fluorescence (AU)
residuals
Time (s)

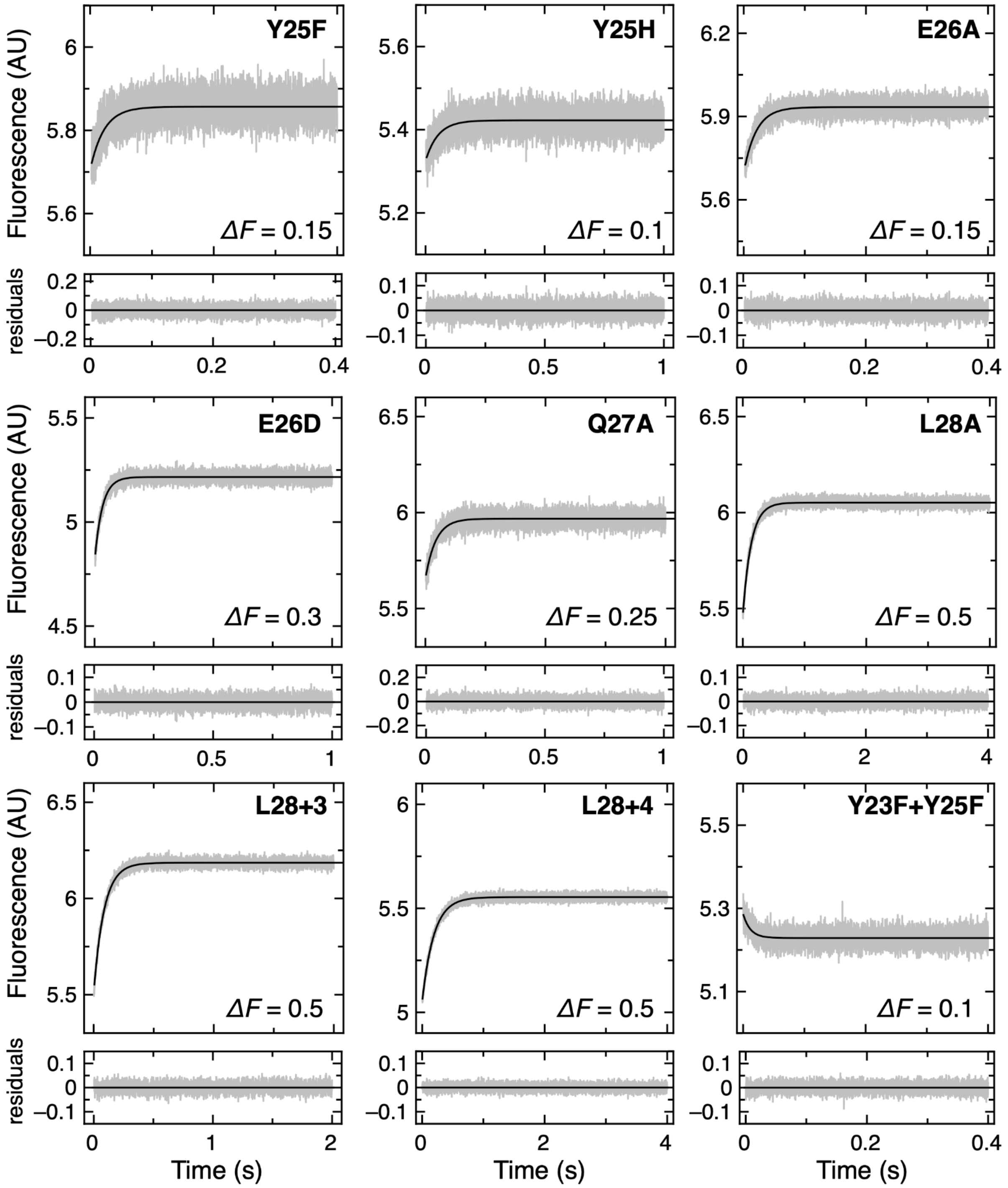


**Figure S1.** ***Association traces for binding of LxCxE mutants to Rb.*** Data were recorded at a constant concentration of FITC-LxCxE peptide after adding different Rb concentrations in pseudo-first order regime. The association trace corresponding to the highest concentration of Rb tested is shown for each mutant. For each mutant, association traces were fit to a mono-exponential function to obtain the observed association rate ($k_{obs}$). The residuals are shown below each plot. The $k_{on}$ value for each mutant was obtained from the slope of the linear fit of $k_{obs}$ versus Rb concentration (see SI Appendix, Supplementary Figure S2, and Supplementary Table S2). $\Delta F$ is the maximal amplitude of the change in fluorescence. The shown traces represent an average of at least 8 individual measurements.

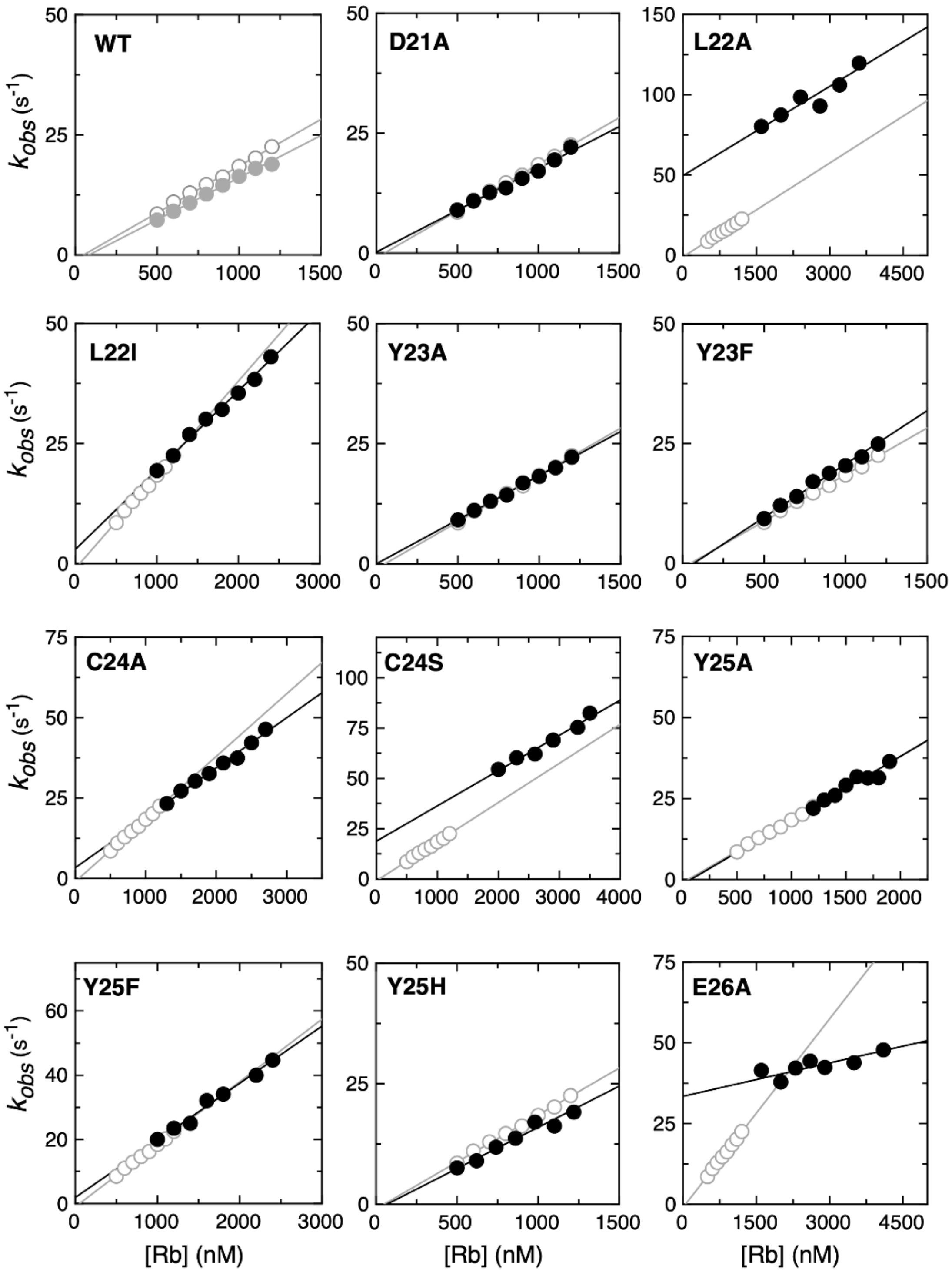
WT
D21A
L22A
L22I
Y23A
Y23F
C24A
C24S
Y25A
Y25F
Y25H
E26A
$k_{obs}$ (s$^{-1}$)
[Rb] (nM)

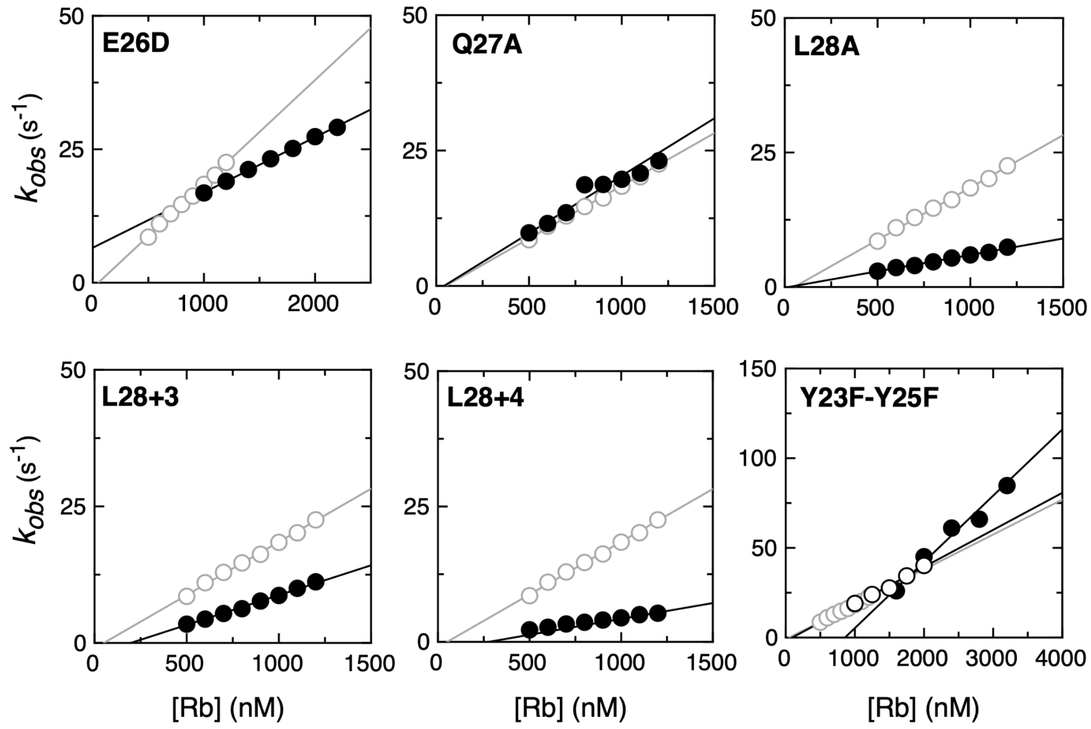


**Figure S2.** ***Pseudo–first-order association plots for LxCxE mutants binding to Rb.*** Association data was recorded at a constant concentration of FITC-LxCxE peptide after adding variable concentrations of Rb (SI Appendix, Supplementary Figure S1). The PFO plots are obtained by plotting the $k_{obs}$ values against Rb concentration (black circles). All PFO plots showed a linear dependence of the $k_{obs}$ on Rb concentration. In every plot except that of the *wild-type* LxCxE peptide, the PFO plot for the *wild-type* LxCxE peptide is shown for comparison (open circles and gray lines). The $k_{on}$ value for each mutant was obtained from the slope of the linear fit of the PFO plot (SI Appendix, Supplementary Table S2). The second PFO plot from the WT (closed gray circles) and Y23-Y25F (open circles and black lines). corresponds to a second replicate done in another laboratory using a different batch of protein and a different stopped-flow apparatus. An error of 5% in the concentration of Rb was considered in the analysis.

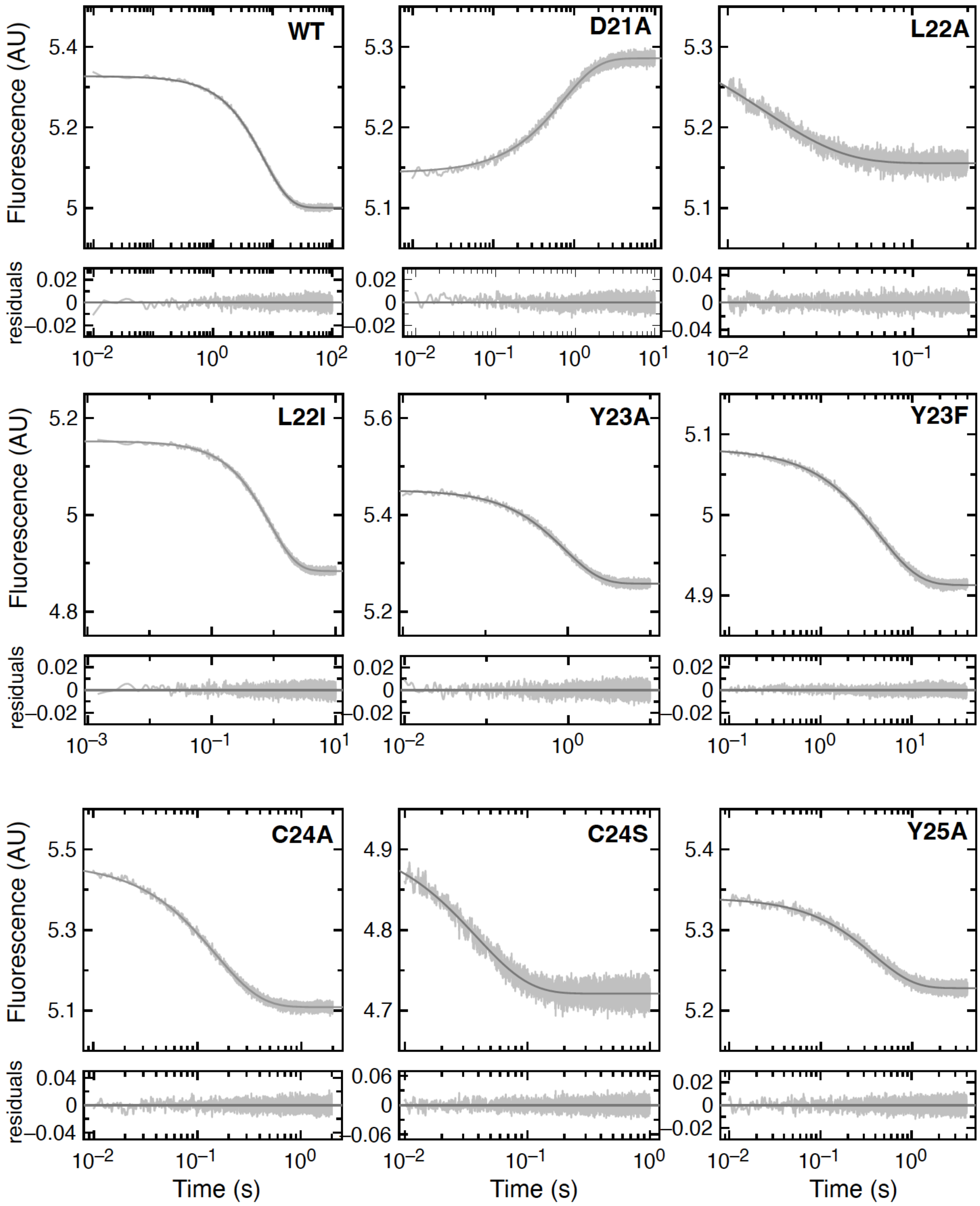

WT
D21A
L22A
L22I
Y23A
Y23F
C24A
C24S
Y25A
Fluorescence (AU)
residuals
Time (s)

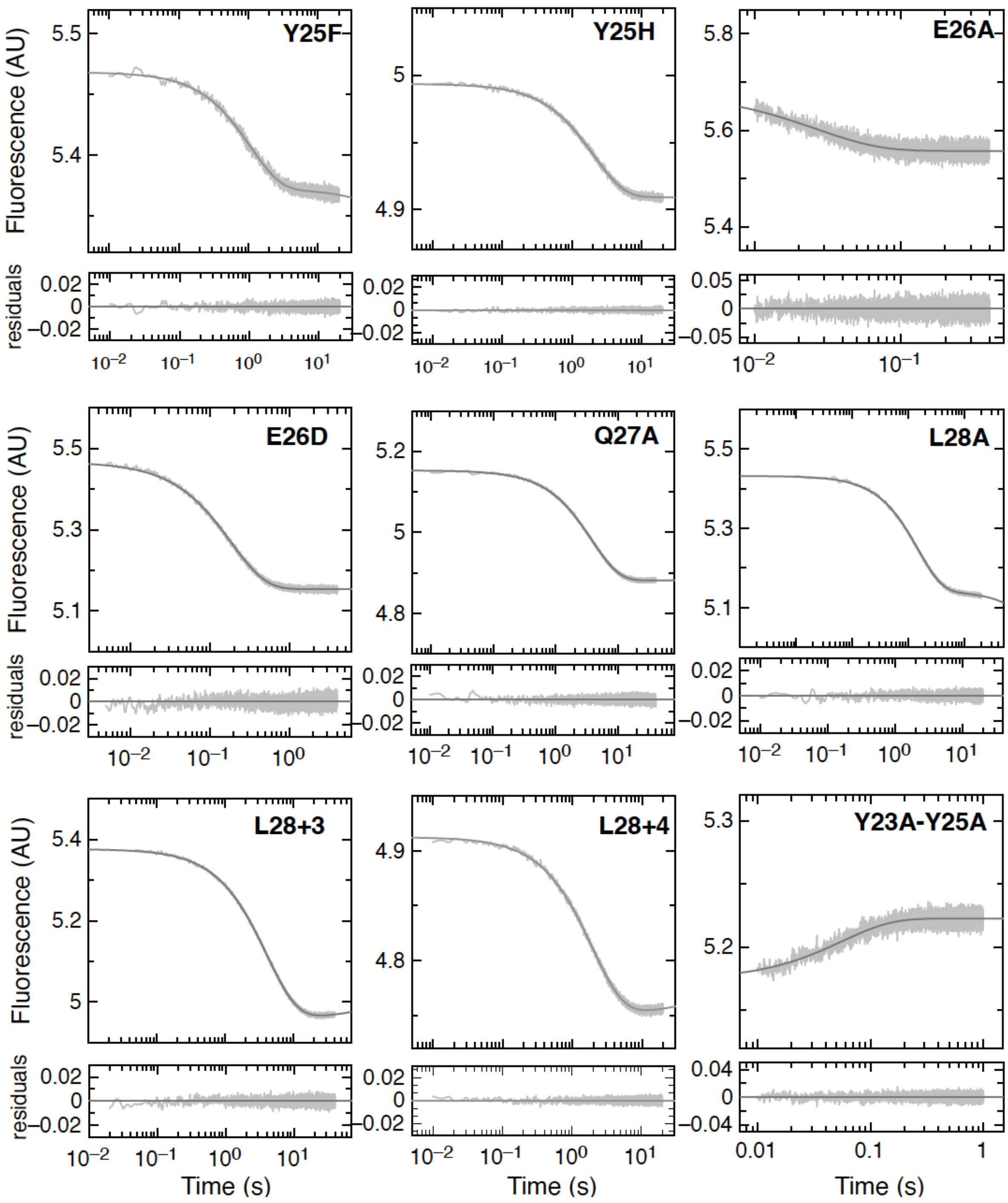

Y25F
Y25H
E26A
E26D
Q27A
L28A
L28+3
L28+4
Y23A-Y25A
Fluorescence (AU)
residuals
Time (s)

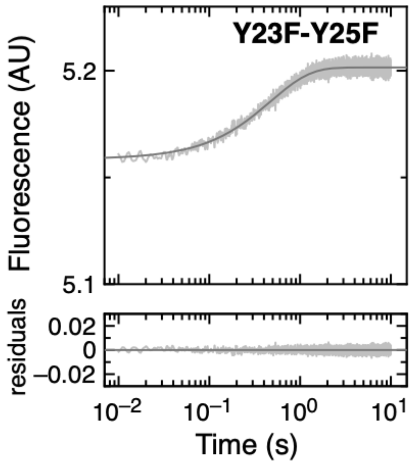


**Figure S3. *Dissociation traces for different LxCxE peptides from Rb.*** The dissociation traces ($k_{off}$) were estimated by displacing a stoichiometric complex of Rb and FITC-labeled peptide with a 20-fold excess of unlabelled peptide. The $k_{off}$ for each mutant were calculated by averaging the rates obtained by fitting the dissociation traces to a mono-exponential decay. For the Y25F, L28A, L28+3 and L28+4 peptides, the dissociation traces were fit to a mono-exponential decay with a linear drift term. In all cases, the residuals are shown below the fit. The $k_{off}$ values for all mutants are reported in SI Appendix, Supplementary Table S2. The shown traces represent an average of at least 8 individual measurements.

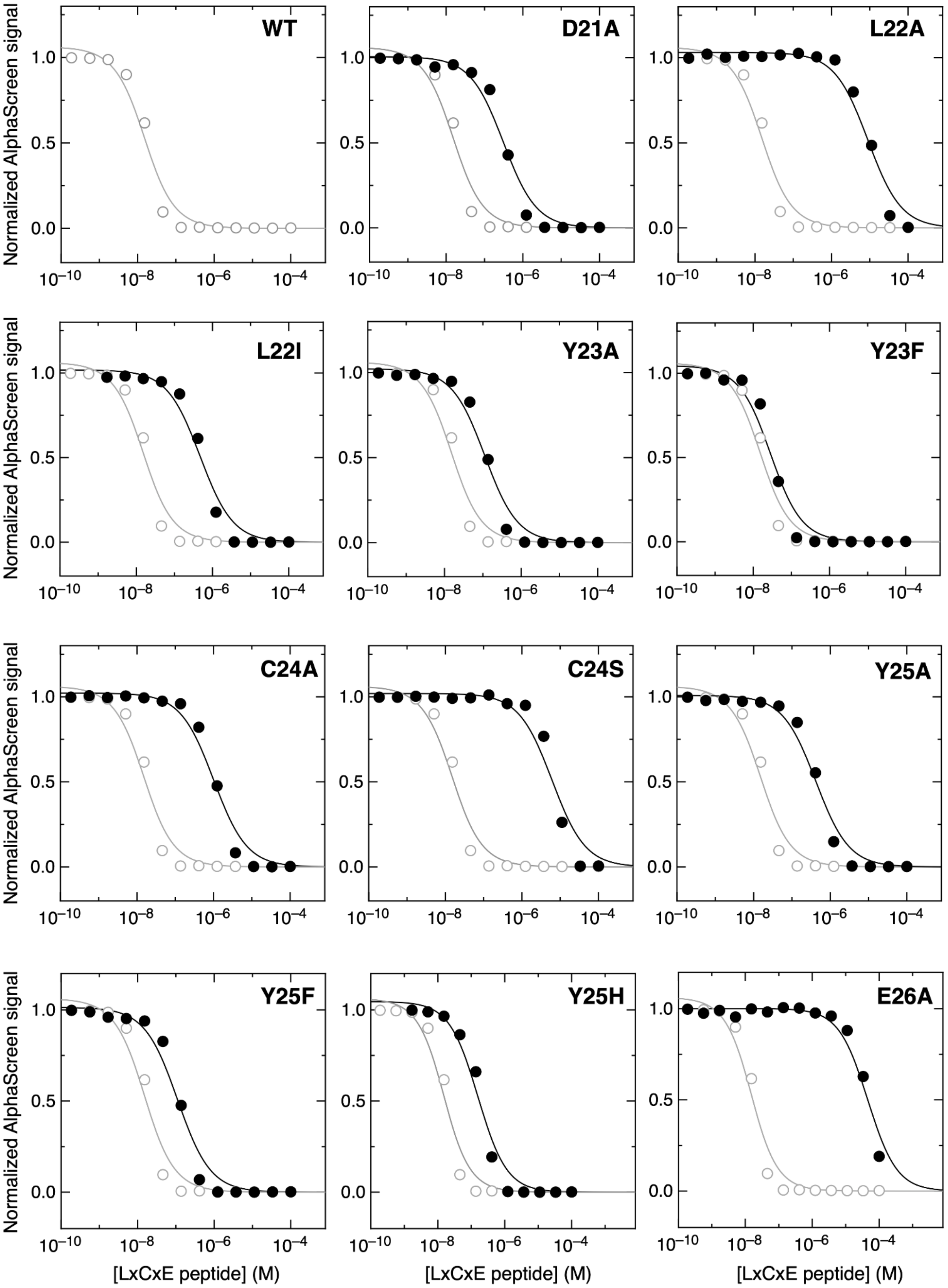
WT
D21A
L22A
L22I
Y23A
Y23F
C24A
C24S
Y25A
Y25F
Y25H
E26A
Normalized AlphaScreen signal
[LxCxE peptide] (M)

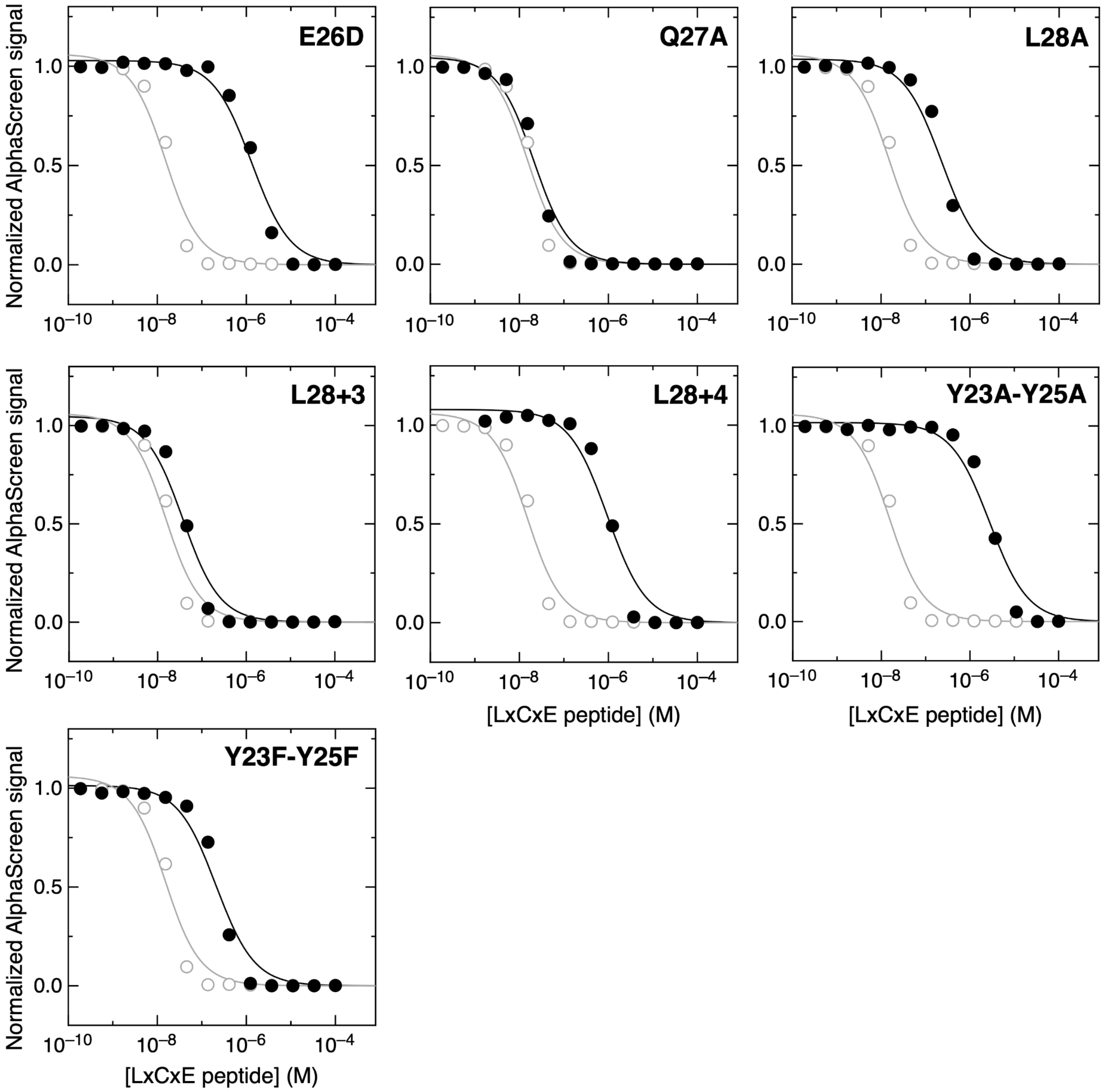


**Figure S4.** ***Alpha Screen affinity measurements of LxCxE peptides for the Rb pocket domain.*** A preformed complex of 6nM His-Rb and 6nM biotin-E7 peptide was competed with increasing concentrations of different LxCxE peptides. In every plot except that of the *wild-type* LxCxE peptide, the plot for the *wild-type* LxCxE peptide is shown for comparison (open circles and gray lines). The affinities of the different peptides are reported in SI Appendix, Table S2.

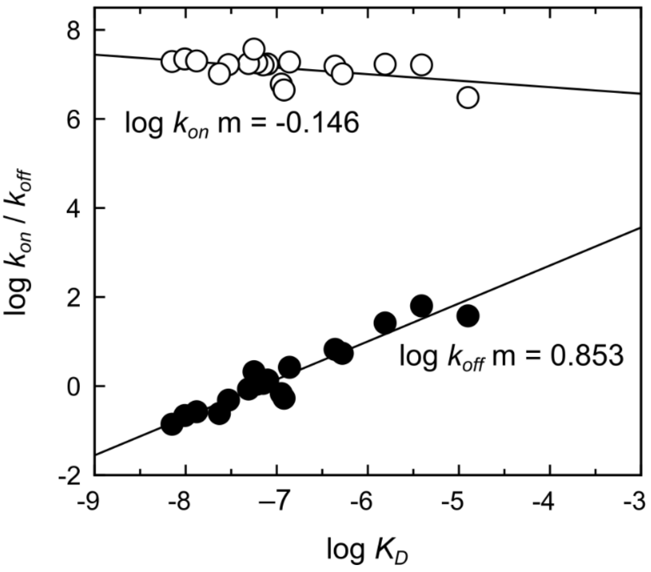


**Figure S5.** ***Linear free energy relationship (LFER) plot for binding of the LxCxE motif to Rb***. The plot shows the dependence of log $k_{on}$ (open circles) and log $k_{off}$ (closed circles) on log $K_D$ for all LxCxE motif mutants. At a global level, changes in $K_D$ are mainly explained by changes in $k_{off}$.

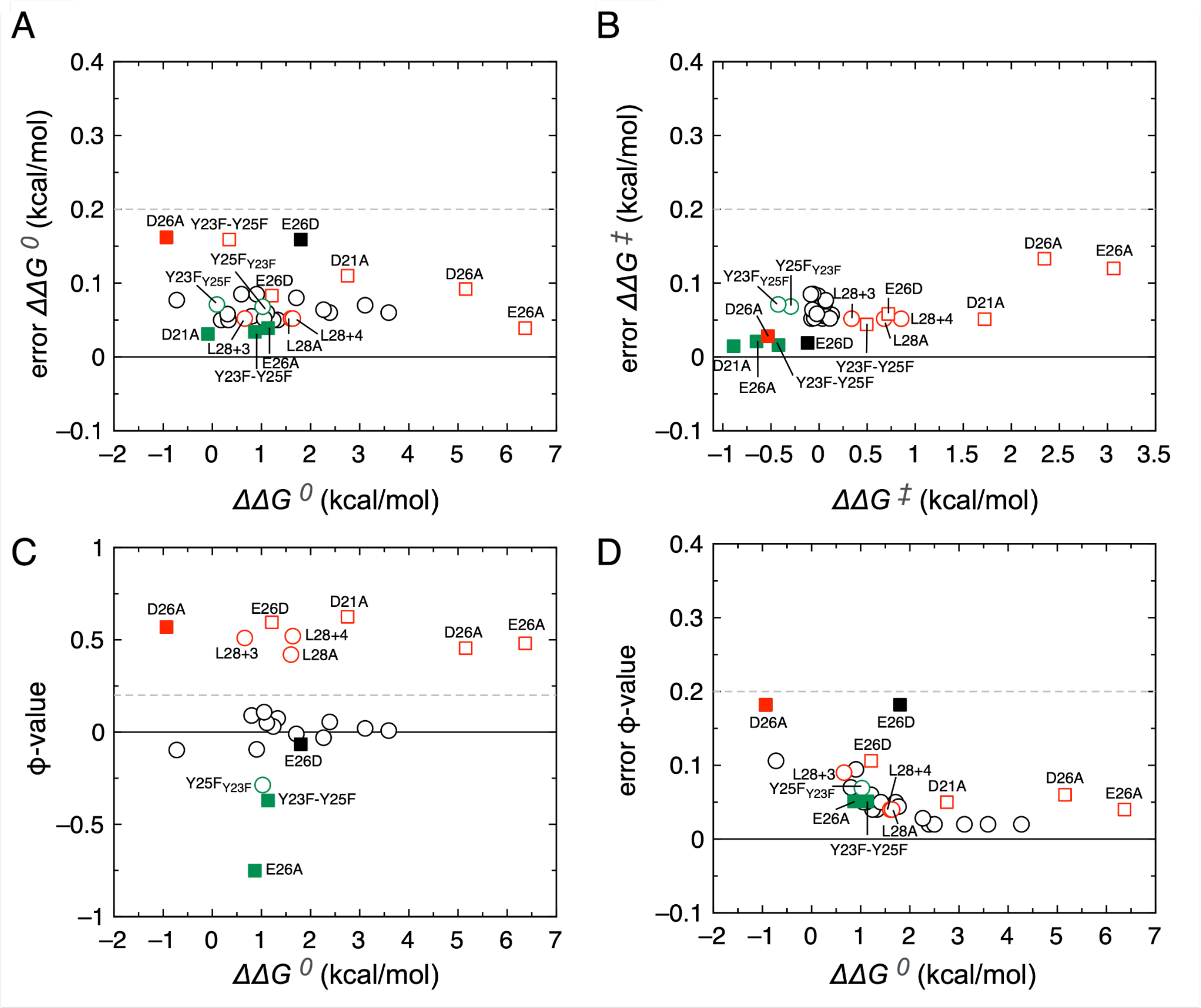


**Figure S6.** ***Correlation of*** $\Delta\Delta G^0$ ***and*** $\Delta\Delta G^\ddagger$ ***values to the errors in the parameters.*** (**A**) Plot of the error (standard deviation) for the change in free energy of binding $\Delta\Delta G^0$ as a function of $\Delta\Delta G^0$. (**B**) Plot of the error (standard deviation) for the change in free energy of binding for the transition state $\Delta\Delta G^\ddagger$ as a function of $\Delta\Delta G^\ddagger$. (**C**) Plot of Φ-values as a function of the change in free energy of binding $\Delta\Delta G^0$. (**D**) Plot of the error of the Φ-values as a function of the change in free energy of binding $\Delta\Delta G^0$ for all mutants. For an assessment of Φ-value uncertainty see Materials and Methods. Empty black circles: mutations producing minor effects on the energetics of the TSE. Squares: Free energy changes of mutants analyzed using a Debye-Hückel approximation to estimate the electrostatic (empty squares) and neutral (filled squares) components. Red: mutations with an intermediate destabilizing effect in the transition state ensemble ($\Delta\Delta G^\ddagger > 0.3$ kcal/mol*); Green:* mutations with opposite effects in the TSE and the final complex (destabilizing mutations with $\Delta\Delta G^\ddagger < 0$ kcal/mol or stabilizing mutations with $\Delta\Delta G^\ddagger > 0$ kcal/mol*)*. An estimated 5% error in the concentration of Rb was considered, and the Debye-Hückel parameter *a* was fixed at 6 Å. All the values are reported in SI Appendix, Table S2 and S11.

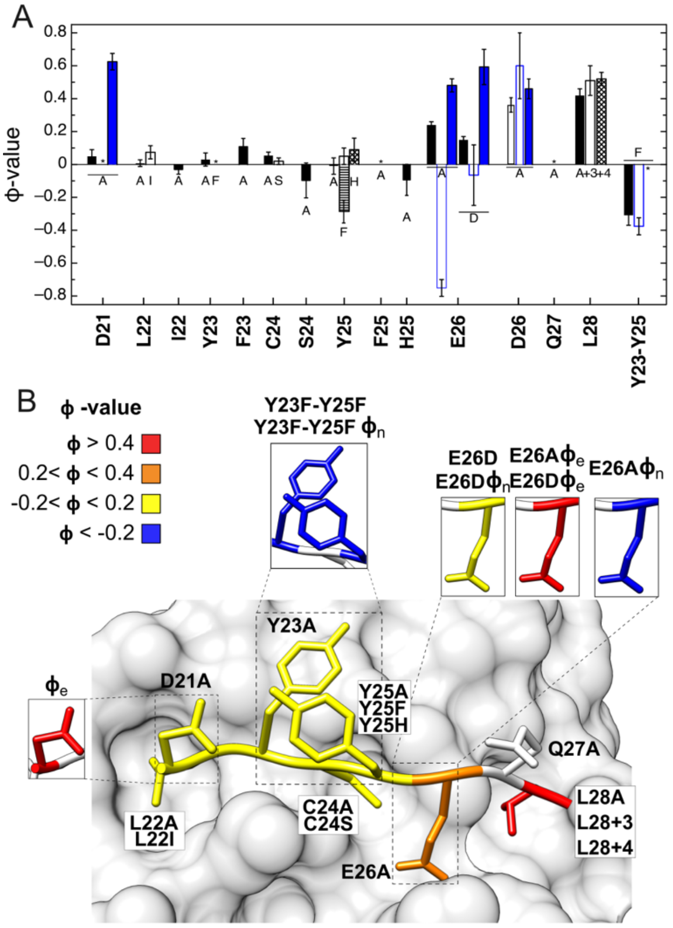


**Figure S7.** ***Analysis of the transition state ensemble for binding of the LxCxE motif to Rb.*** (**A**) Individual Φ-values obtained for all LxCxE motif mutations. The Φ-values corresponding to mutations at each position are depicted as bars. Black bars: mutations to Alanine; White bars: conservative substitutions (L22I, Y23F, C24S, Y25F, Y23F-Y25F, E26D, L28+3); Diagonal striped bars: Y25H, L28+4. Horizontal striped bar: Y25F mutation performed on the F23 background F23-Y25 → F23-F25 ($Y25F_{F23}$). The Φ-values for LxCxE charge mutants are reported as electrostatic ($\Phi_e$: Closed blue bars) and neutral ($\Phi_n$: Open blue bars) components obtained from the ionic strength dependence of association and dissociation rate constants using the Debye-Hückel approximation. (**B**) Graphic representation of representative Φ-values obtained at different positions of the LxCxE motif. Negative Φ-values are obtained at three different positions (D21, Y23 and Y25).

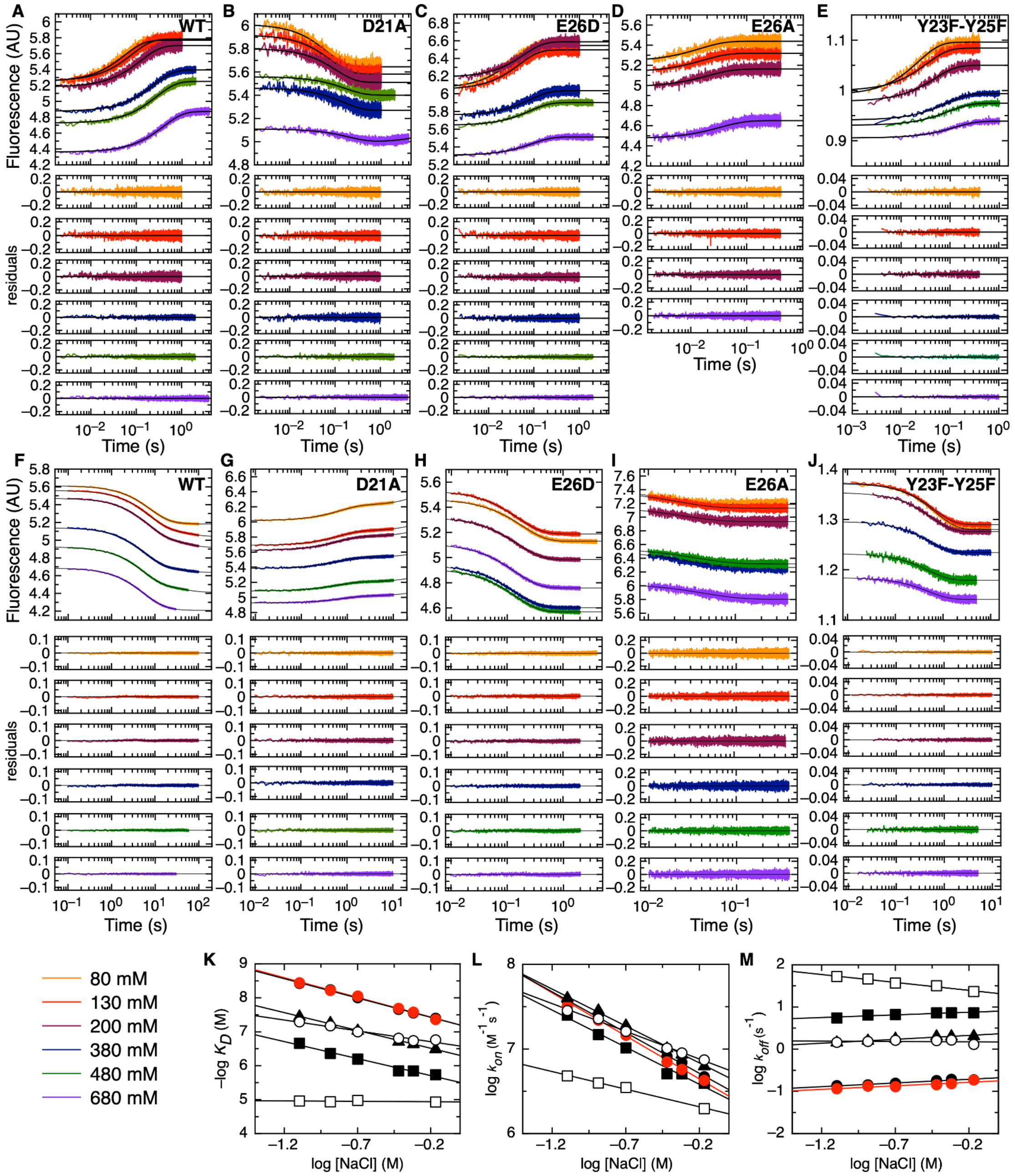


**Figure S8.** ***Kinetic and equilibrium analysis of the interaction between Rb and LxCxE charge mutants at varying NaCl concentrations*** Data were recorded at a constant concentration of FITC-LxCxE peptide and Rb at different NaCl concentrations. For each mutant, association traces were fitted to a mono-exponential function to obtain the observed association rate ($k_{obs}$) (**A-E**). The dissociation rate constants ($k_{off}$) for each mutant were calculated by averaging the rates obtained by fitting the dissociation traces to a mono-exponential decay (**F-J**). In all cases, the residuals are shown below the fit. The shown traces represent an average of at least 8 individual measurements. (**K**) NaCl dependence of the equilibrium dissociation constant ($K_D$). In all cases, the data were fit to linear functions, with the $\Gamma_{EQ}$ -1.14 ± 0.03 and -1.07 ± 0.02 for the WT, -0.64 ± 0.03 for D21A, -0.96 ± 0.04 for E26D, -0.03 ± 0.06 for E26A and -0.93 ± 0.02 for Y23F-Y25F. (**L**) NaCl dependence of the kinetic association rate constant ($k_{on}$). In all cases, the data were fit to linear functions, with the $\Gamma_{ON}$ -0.96 ± 0.03 and -0.92 ± 0.02 for the WT, -0.65 ± 0.03 for D21A, -0.85 ± 0.04 for E26D, -0.51 ± 0.06 for E26A and -0.82 ± 0.02 for Y23F-Y25F. (**M**) NaCl dependence of kinetic dissociation rate constant ($k_{off}$). In all cases, the data were fit to linear functions, with the $\Gamma_{OFF}$ 0.202 ± 0.001 and 0.156 ± 0.001 for the WT, 0.047 ± 0.001 for D21A, 0.112 ± 0.001 for E26D, -0.484 ± 0.006 for E26A and 0.147 ± 0.01 for Y23F-Y25F. Black and red circles: LxCxE$_{WT}$ and its replicate; Open circles: D21A; Closed squares: E26D; Open squares: E26A; Closed triangles: Y23F-Y25F.

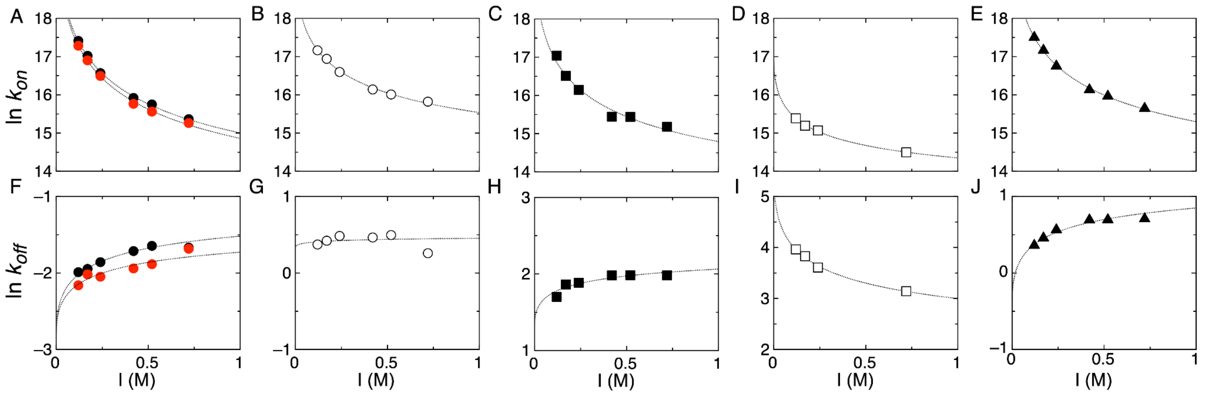


**Figure S9.** ***Analysis of the Ionic strength dependence of the interaction between* LxCxE charge mutants and Rb using the *Debye-Hückel model.*** (**A-E**) Ionic strength dependence of the association rate constants. (**F-J**). Ionic strength dependence of the dissociation rate constants. Data from the plots were globally fitted for all mutants using Eq. (6) and the values of ln $k_{on,basal}$ and $U_{on}$/RT or those of ln $k_{off,basal}$ and $U_{off}$/RT were obtained as fitting parameters for each mutant (see Materials and Methods, SI Appendix, Table S11). The constant b was kept fixed using a value for the *a* parameter of 6 Å. Black and red circles: WT and its replicate; Open circles: D21A; Closed squares: E26D; Open squares: E26A and Closed triangles: Y23F-Y25F.

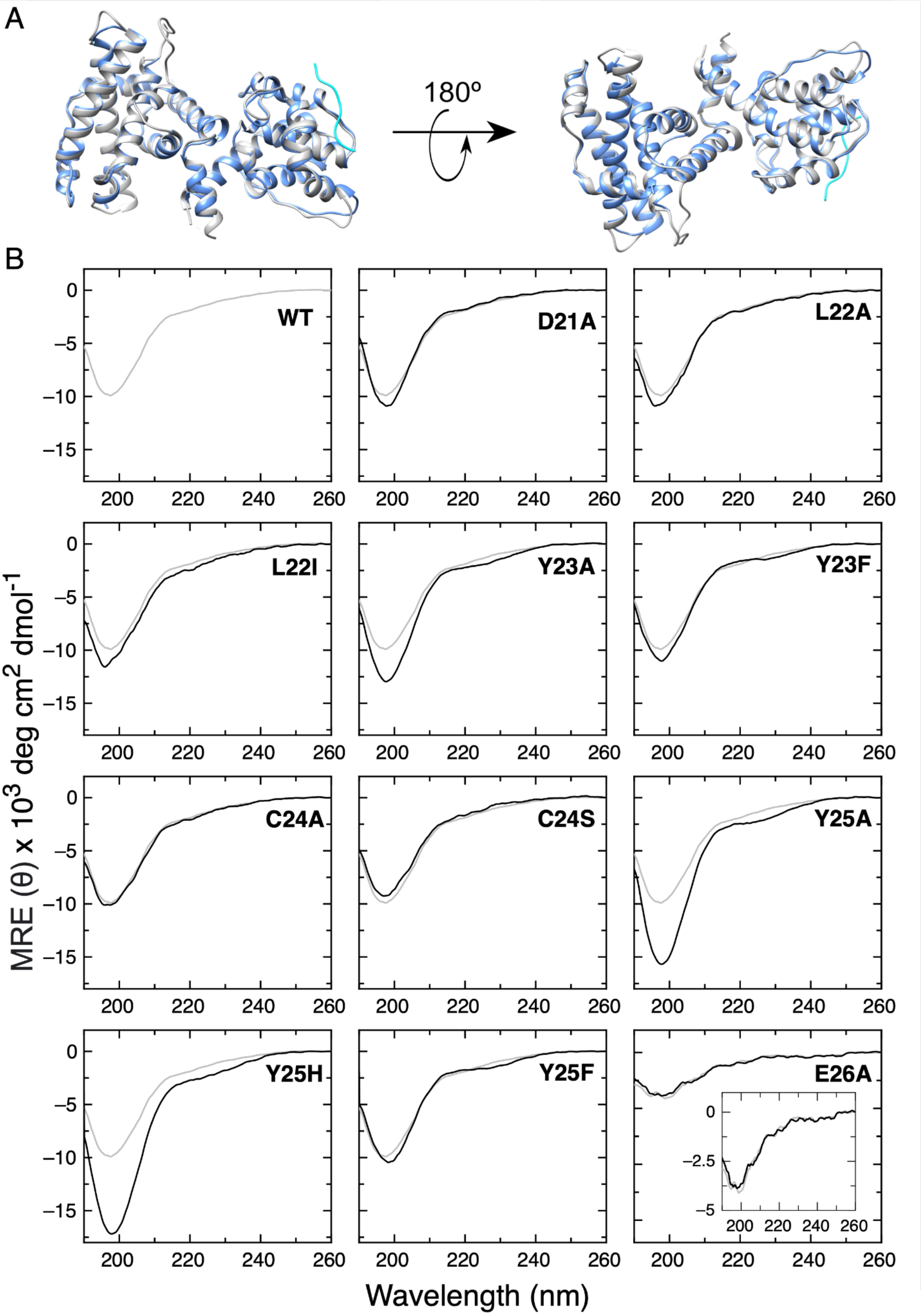
A
180º
B
WT
D21A
L22A
L22I
Y23A
Y23F
C24A
C24S
Y25A
Y25H
Y25F
E26A
MRE (θ) x 10³ deg cm² dmol⁻¹
Wavelength (nm)
0
−5
−10
−15
200
220
240
260
−2.5

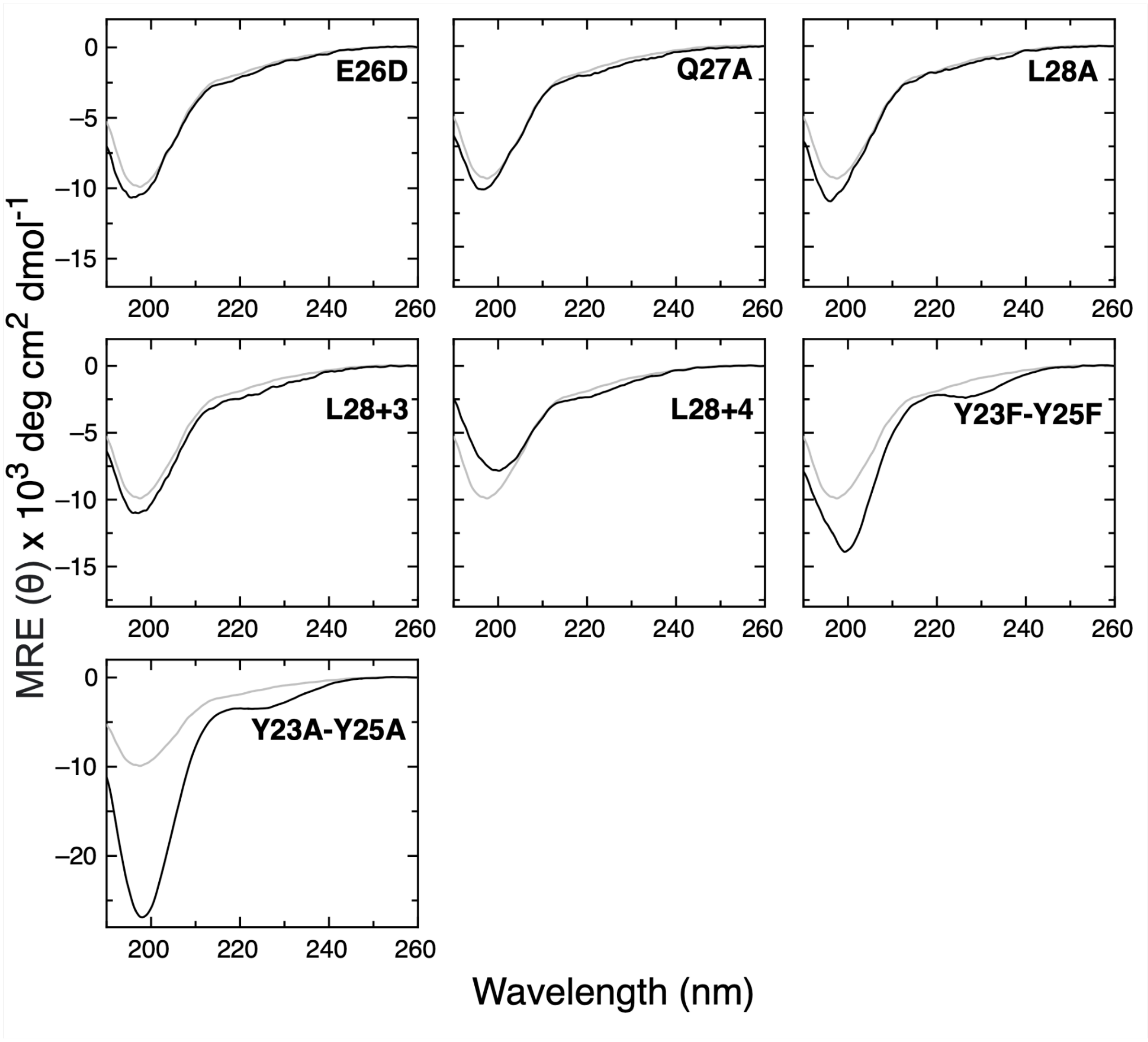


**Figure S10. (A)** ***Overlay of the crystal structures of the unliganded RbAB domain and the RbAB domain bound to the E7 LxCxE motif.*** Cartoon representation of the unligated Rb domain (gray, PDB ID: 3POM) aligned against the Rb domain in complex with the HPV E7 21-29 peptide containing the LxCxE motif (PDB ID: 1GUX, cyan). The LxCxE motif is shown in dark blue. The global RMSD value of the alignment is 0.526 Å. This indicates that there are no significant changes in the structure of the Rb domain when the complex is formed, suggesting that the Rb domain behaves as a fairly rigid receptor when binding to LxCxE motifs. (**B)** Far-UV CD spectra of acetylated LxCxE motif peptides including $LxCxE_{WT}$ (gray line) and all mutants (black line). Measurements were performed in 20 mM sodium phosphate buffer, 1mM DTT and pH 7.0 at 20 ºC. In every plot except that of $LxCxE_{WT}$, the CD spectrum for $LxCxE_{WT}$ is shown for comparison, except for the E26A mutant. For E26A, the spectrum corresponds to the FITC-labeled $LxCxE_{E26A}$ peptide compared to the FITC-labeled $LxCxE_{WT}$ peptide, where both spectra have excellent overlap but the minimum is less pronounced due to the absorbance of the fluorophore.

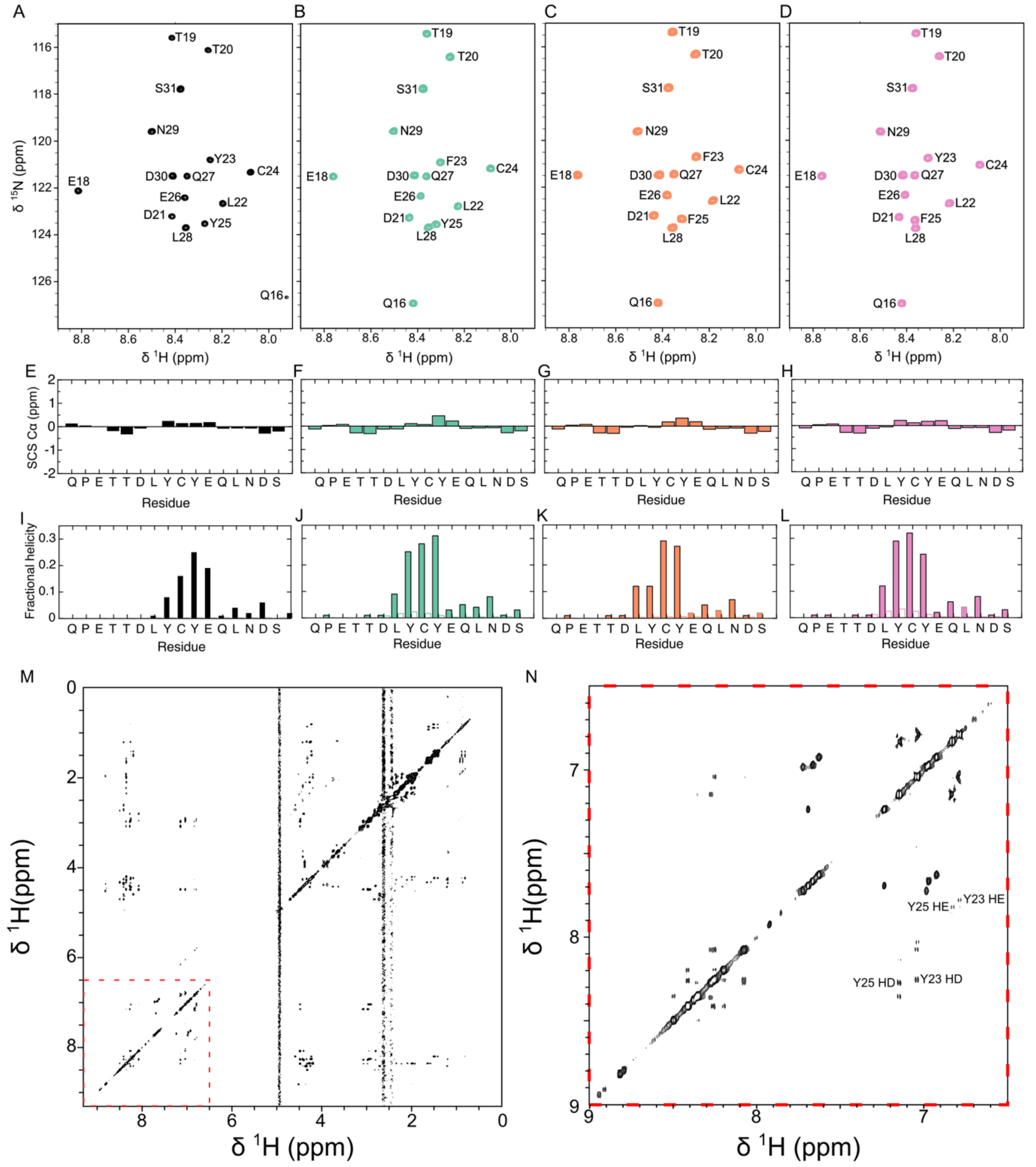


**Figure S11***. Secondary structure analysis of the LxCxE peptide variants with NMR.* (**A-D**) $^{1}$H, $^{15}$N HSQC spectra of LxCxE$_{WT}$ (black), LxCxE$_{Y23F}$ (green), LxCxE$_{Y25F}$ (orange) and LxCxE$_{Y23F\text{-}Y25F}$ (pink) and their assigments. (**E-H**) Secondary Cα subindices chemical shifts. (**I-L**) Secondary structure analysis of LxCxE peptide variants with NMR data processed by CSI calculations (closed bars) or by the δ2Δ method (empty bars). For both, fractional helicity =1 corresponds to 100% helicity. (**M-N**) Full $^{1}$H, $^{1}$H-2D NOESY of LxCxE$_{WT}$ (**M**) and inset showing the region corresponding to resonances for Y23/Y25 (red dashes in M and expanded in panel **N**). NOEs between aromatic ring protons of the tyrosines are absent, indicating that there is no observable interaction between Y23 and Y25 in the free peptide.

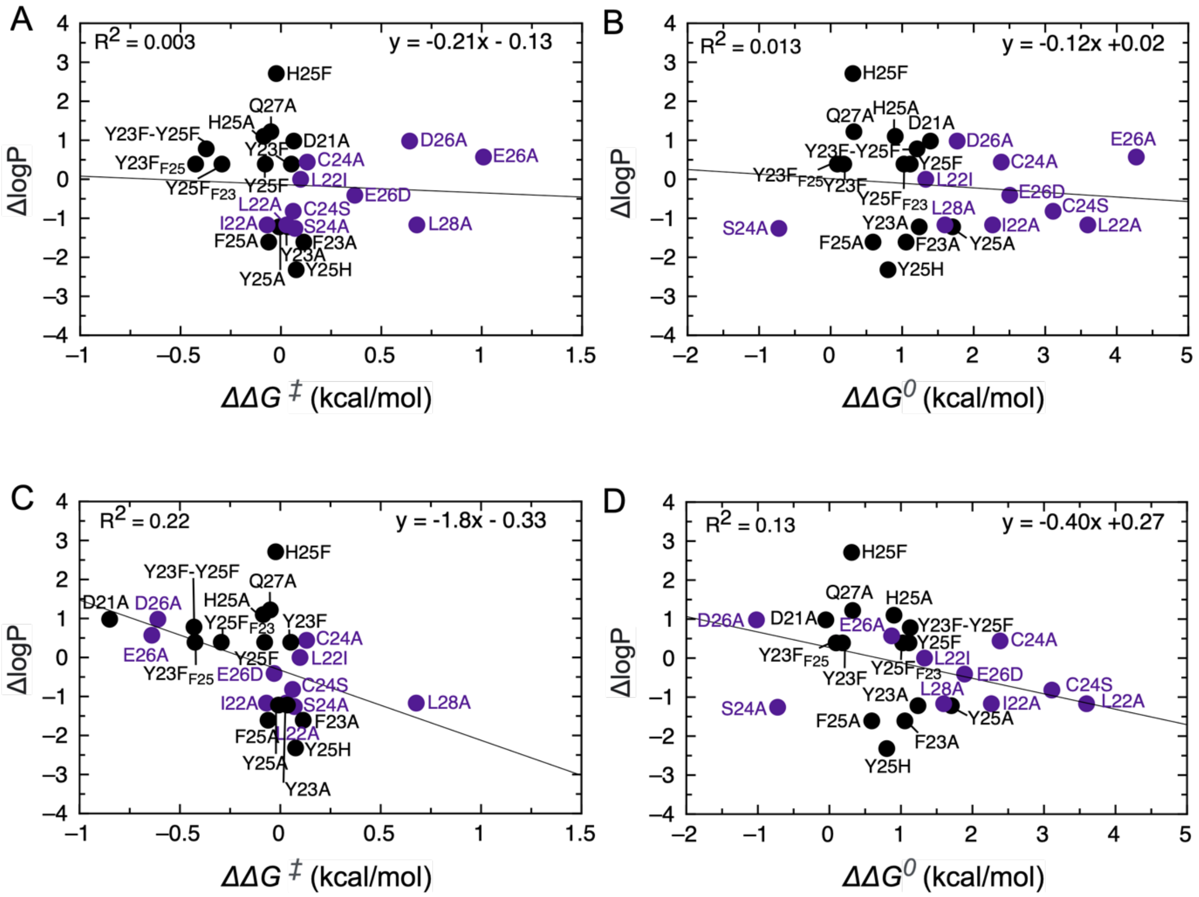


***Figure S12. Correlation between changes in the hydrophobicity of LxCxE motif mutants and changes in the transition state ensemble or equilibrium free energies of binding.*** The change in hydrophobicity for each of the LxCxE motif mutants (ΔlogP) is plotted against (**A,C**) the difference in free energy of binding for the transition state ($\Delta\Delta G^{\ddagger}$) or (**B,D**) the difference in the free energy of binding at equilibrium ($\Delta\Delta G^{0}$). ΔlogP (n-octanol/water) was calculated using ChemDraw, and a more detailed description of the calculation of the change in hydrophobicity is provided in Materials and Methods. In both plots, contact positions are shown as purple circles and non-contact positions are shown as black circles. The global linear fit of the data is represented in black and the values obtained are y = -0.21x - 0.13; $R^2$ = 0.003 for $\Delta\Delta G^{\ddagger}$ (**A**) and y = -0.12x + 0.02; $R^2$ = 0.013 for $\Delta\Delta G^{0}$ (**B**). Both plots reveal no significant correlation between free energy of binding and changes in hydrophobicity. Even when considering $\Delta\Delta G^{\ddagger}$ values originating from the non-electrostatic component for D21A, E26A, E26D, D26A and Y23F-Y25F this does not change the analysis with a slope value of -1.8 and $R^2$ =0.22 for the dependence of $\Delta\Delta G^{\ddagger}$ on ΔlogP (**C**) and y = -0.40x + 0.27; $R^2$ = 0.13 for $\Delta\Delta G^{0}$ (**D**) For a discussion of changes in hydrophobicity see SI Appendix, Supplementary Text.

**Table S1.** ***Sequence of the LxCxE motif variants used for kinetic studies.***

| Peptide | Sequence |
|---|---|
| **WT** | QPETTDLYCYEQLNDS |
| **D21A** | QPETTALYCYEQLNDS |
| **L22A** | QPETTDAYCYEQLNDS |
| **L22I** | QPETTDIYCYEQLNDS |
| **Y23A** | QPETTDLACYEQLNDS |
| **Y23F** | QPETTDLFCYEQLNDS |
| **C24A** | QPETTDLYAYEQLNDS |
| **C24S** | QPETTDLYSYEQLNDS |
| **Y25A** | QPETTDLYCAEQLNDS |
| **Y25F** | QPETTDLYCFEQLNDS |
| **Y25H** | QPETTDLYCHEQLNDS |
| **E26A** | QPETTDLYCYAQLNDS |
| **E26D** | QPETTDLYCYDQLNDS |
| **Q27A** | QPETTDLYCYEALNDS |
| **L28A** | QPETTDLYCYEQANDS |
| **L28+3** | QPETTDLYCYEQALND |
| **L28+4** | QPETTDLYCYEQAALN |
| **Y23A-Y25A** | QPETTDLACAEQLNDS |
| **Y23F-Y25F** | QPETTDLFCFEQLNDS |

*Red font: mutated residues*

**Table S2.** ***Kinetic rate constants and thermodynamic data for the binding of wild-type and mutant LxCxE motif considering an error in [Rb] of 5%.***

| | $k_{on}$ x $10^7$ ($M^{-1}$ $s^{-1}$) | $k_{off}$ ($s^{-1}$) | $K_{D,\,KIN}$ (nM)# | %$K_{D,\,EQ}$ (nM) | $\Delta G_{KIN}$ (kcal/mol)* | $\Delta\Delta G^0$ (kcal/mol)$ | $\Delta\Delta G^‡$ (kcal/mol)^ | ϕ-value |
|---|---|---|---|---|---|---|---|---|
| **WT**** | 1.95 ± 0.12<br>1.76 ± 0.11 | 0.139 ± 0.001<br>0.126 ± 0.001 | 7.14 ± 0.45<br>7.19 ± 0.47 | 8 ± 2 | -10.93 ± 0.04<br>-10.92 ± 0.04 | 0<br>0 | 0<br>0 | -<br>- |
| **D21A** | 1.75 ± 0.11 | 1.366 ± 0.003 | 78 ± 5 | 273 ± 58 | -9.53 ± 0.04 | 1.39 ± 0.05 | 0.06 ± 0.05 | 0.05 ± 0.04 |
| **L22A** | 1.85 ± 0.16 | 63.3 ± 0.3 | 3419 ± 303 | 8814 ± 1529 | -7.33 ± 0.05 | 3.59 ± 0.06 | 0.03 ± 0.06 | 0.01 ± 0.02 |
| **L22I** | 1.65 ± 0.11 | 1.15 ± 0.001 | 70 ± 5 | 463 ± 93 | -9.60 ± 0.04 | 1.33 ± 0.05 | 0.10 ± 0.05 | 0.07 ± 0.04 |
| **Y23A** | 1.83 ± 0.12 | 1.100 ± 0.002 | 60 ± 4 | 94 ± 20 | -9.69 ± 0.04 | 1.24 ± 0.05 | 0.035 ± 0.05 | 0.03 ± 0.04 |
| **Y23F** | 2.23 ± 0.14 | 0.218 ± 0.001 | 9.8 ± 0.6 | 22 ± 6 | -10.74 ± 0.04 | 0.18 ± 0.05 | -0.08 ± 0.05 | -† |
| **C24A** | 1.56 ± 0.12 | 6.680 ± 0.009 | 430 ± 32 | 835 ± 169 | -8.54 ± 0.04 | 2.39 ± 0.06 | 0.131 ± 0.06 | 0.06 ± 0.02 |
| **C24S** | 1.75 ± 0.19 | 26.1 ± 0.1 | 1489 ± 161 | 5765 ± 1263 | -7.82 ± 0.06 | 3.11 ± 0.07 | 0.06 ± 0.07 | 0.02 ± 0.02 |
| **Y25A** | 1.98 ± 0.25 | 2.639 ± 0.006 | 133 ± 17 | 357 ± 79 | -9.22 ± 0.07 | 1.71 ± 0.08 | -0.01 ± 0.08 | -0.01 ± 0.05 |
| **Y25F** | 1.78 ± 0.13 | 0.860 ± 0.002 | 48 ± 3 | 88 ± 19 | -9.81 ± 0.04 | 1.11 ± 0.06 | 0.05 ± 0.06 | 0.05 ± 0.05 |
| **Y25H** | 1.71 ± 0.12 | 0.484 ± 0.001 | 28 ± 2 | 155 ± 31 | -10.12 ± 0.04 | 0.80 ± 0.06 | 0.08 ± 0.05 | 0.09 ± 0.07 |
| **E26A** | 0.34 ± 0.04 | 37.8 ± 0.5 | 10955 ± 1181 | 38727 ± 8054 | -6.65 ± 0.06 | 4.27 ± 0.07 | 1.01 ± 0.07 | 0.24 ± 0.02 |
| **E26D** | 1.04 ± 0.07 | 5.446 ± 0.006 | 526 ± 33 | 1145 ± 250 | -8.42 ± 0.04 | 2.50 ± 0.05 | 0.37 ± 0.05 | 0.15 ± 0.02 |
| **Q27A** | 2.12 ± 0.14 | 0.265 ± 0.001 | 12.5 ± 0.8 | 13 ± 3 | -10.60 ± 0.04 | 0.33 ± 0.05 | -0.05 ± 0.05 | -† |
| **L28A** | 0.61 ± 0.04 | 0.677 ± 0.001 | 111 ± 7 | 202 ± 49 | -9.33 ± 0.04 | 1.60 ± 0.05 | 0.68 ± 0.05 | 0.42 ± 0.04 |
| **L28+3** | 1.09 ± 0.07 | 0.242 ± 0.001 | 22 ± 1 | 29 ± 7 | -10.27 ± 0.04 | 0.66 ± 0.05 | 0.34 ± 0.05 | 0.51 ± 0.09 |
| **L28+4** | 0.45 ± 0.03 | 0.534 ± 0.001 | 119 ± 8 | 942 ± 229 | -9.29 ± 0.04 | 1.64 ± 0.05 | 0.85 ± 0.05 | 0.52 ± 0.04 |
| **Y23A-Y25A** | - | 17.7 ± 0.1 | - | 2309 ± 431 | - | 2.75 ± 0.20 | - | - |
| **Y23F-Y25F**** | 3.7 ± 0.4<br>2.1 ± 0.2 | 2.10 ± 0.008<br>1.68 ± 0.003 | 57 ± 6<br>82 ± 8 | 196 ± 41 | -9.72 ± 0.06<br>-9.51 ± 0.06 | 1.21 ± 0.07<br>1.41 ± 0.07 | -0.37 ± 0.07<br>-0.09 ± 0.07 | -0.31 ± 0.06<br>-0.07 ± 0.05 |
| **I22A** | - | - | - | - | - | 2.26 ± 0.06 | -0.069 ± 0.06 | -0.03 ± 0.03 |
| **S24A** | - | - | - | - | - | -0.72 ± 0.08 | 0.07 ± 0.08 | -0.1 ± 0.1 |
| **D26A** | - | - | - | - | - | 1.77 ± 0.07 | 0.64 ± 0.07 | 0.36 ± 0.04 |
| **Y23F$_{F25}$** | - | - | - | - | - | 0.09 ± 0.07 | -0.43 ± 0.07 | -† |
| **Y25F$_{F23}$** | - | - | - | - | - | 1.03 ± 0.07 | -0.29 ± 0.07 | -0.29 ± 0.07 |
| **F23A** | - | - | - | - | - | 1.06 ± 0.05 | 0.11 ± 0.05 | 0.11 ± 0.05 |
| **F25A** | - | - | - | - | - | 0.59 ± 0.09 | -0.06 ± 0.09 | -† |
| **H25A** | - | - | - | - | - | 0.90 ± 0.09 | -0.09 ± 0.08 | -0.09 ± 0.09 |
| **H25F** | - | - | - | - | - | 0.31 ± 0.06 | -0.02 ± 0.06 | -† |

The errors for $k_{on}$ were obtained from the standard deviation of fitted parameters of the pseudo-first-order association kinetics considering an error of Rb of 5%. Errors for $k_{off}$ were obtained from the fit of the average of all measured dissociation traces (Figure 1 and SI Appendix, Figures SI 1-3). # $K_{D,\,KIN} = k_{off}/k_{on}$. % $K_{D,\,EQ}$ were calculated from competition experiments performed using an AlphaScreen assay (see Materials and Methods section). *$\Delta G_{kin}=RT\ ln\ (K_{D\ kin})$. $ $\Delta\Delta G^0 = \Delta G^0_{MUT} - \Delta G^0_{WT}$. ^$\Delta\Delta G^‡ = \Delta G^‡_{MUT} - \Delta G^‡_{WT}$. ** The second value corresponds to a second replicate done in another laboratory using a different batch of protein and a different stopped-flow apparatus. Y23F$_{F25}$ and Y25F$_{F23}$ correspond to the Y23-F25 → F23-F25 and F23-Y25 → F23-F25 mutations respectively. †We do not report these Φ-values because the $\Delta\Delta G^0$ value is lower than 0.6 kcal/mol. Errors for $K_{D,\,KIN}$, $\Delta G_{KIN}$, $\Delta\Delta G^0$, $\Delta\Delta G^‡$ and for Φ-values were obtained from the primary parameters using standard equations for error propagation.

**Table S3.** ***Experimental conditions for measuring the association and dissociation kinetics of wild-type and mutant LxCxE motifs to Rb.***

| | **Association reactions** | | | | | **Dissociation reactions** | | | | **$t_{1/2}$ (s)** |
|---|---|---|---|---|---|---|---|---|---|---|
| | **[Rb] (nM)** | **[FITC-peptide] (nM)** | **TB (s)*** | **f(complex)**** | **traces** | **[Complex] (nM)** | **[peptide] (nM)** | **TB (s)*** | **f(complex)**** | |
| **WT** | 500-1200 | 100 | 1 | 0.98-0.99 | > 7 | 100 | 2000 | 100 | 0.77 | 4.983 ± 0.003<br>#5.50 ± 0.01 |
| **D21A** | 500-1200 | 100 | 1 | 0.85-0.93 | 9 | 500 | 10000 | 10 | 0.67 | 0.507 ± 0.001 |
| **L22A** | 1600-3600 | 400 | 0.2 | 0.27-0.46 | > 11 | 6000 | 60000 | 0.2 | 0.45 | 0.011 ± 0.001 |
| **L22I** | 1000-2400 | 200 | 1 | 0.92-0.97 | > 8 | 2000 | 40000 | 10 | 0.83 | 0.601 ± 0.001 |
| **Y23A** | 500-1200 | 100 | 1 | 0.87-0.95 | 9 | 500 | 10000 | 10 | 0.77 | 0.632 ± 0.001 |
| **Y23F** | 500-1200 | 100 | 1 | 0.98-0.99 | > 8 | 500 | 10000 | 40 | 0.61 | 3.182 ± 0.005 |
| **C24A** | 1300-2700 | 250 | 0.4 | 0.72-0.85 | > 8 | 913 | 18000 | 2 | 0.51 | 0.104 ± 0.001 |
| **C24S** | 2340-3159 | 400 | 0.2 | 0.5-0.58 | > 7 | 1700 | 40000 | 1 | 0.34 | 0.026 ± 0.001 |
| **Y25A** | 1200-1900 | 200 | 0.4 | 0.88-0.93 | >6 | 2000 | 40000 | 4 | 0.77 | 0.263 ± 0.001 |
| **Y25F** | 1000-2400 | 200 | 1 | 0.87-0.96 | >7 | 500 | 10000 | 20 | 0.7 | 0.806 ± 0.002 |
| **Y25H** | 500-1220 | 100 | 1 | 0.93-0.97 | >8 | 2000 | 40000 | 20 | 0.89 | 1.431 ± 0.001 |
| **E26A** | 1600-4100 | 400 | 0.4 | 0.12-0.22 | > 8 | 2000 | 40000 | 0.4 | 0.11 | 0.018 ± 0.001 |
| **E26D** | 1000-2400 | 200 | 1 | 0.62-0.81 | > 7 | 2000 | 40000 | 4 | 0.60 | 0.127 ± 0.001 |
| **Q27A** | 500-1200 | 100 | 1 | 0.97-0.99 | >8 | 500 | 10000 | 40 | 0.85 | 2.615 ± 0.001 |
| **L28A** | 500-1200 | 100 | 4 | 0.78-0.9 | >7 | 500 | 10000 | 20 | 0.62 | 1.024 ± 0.001 |
| **L28+3** | 500-1200 | 100 | 2 | 0.95-0.98 | >8 | 500 | 10000 | 40 | 0.81 | 2.861 ± 0.002 |
| **L28+4** | 500-1200 | 100 | 4 | 0.78-0.9 | >7 | 500 | 5000 | 20 | 0.62 | 1.297 ± 0.010 |
| **Y23A-Y25A** | - | - | - | - | - | 4000 | 80000 | 1 | 0.61 | 0.039 ± 0.001 |
| **Y23F-Y25F#** | 1600-3200<br>1000-2400 | 400<br>200 | 0.4<br>0.4 | 0.95-0.98<br>0.91-0.96 | >10<br>>12 | 3000<br>1000 | 60000<br>20000 | 20<br>10 | 0.87<br>0.82 | 0.331 ± 0.001<br>0.412 ± 0.001 |

*TB refers to the time base used for the trace recording in each experiment. **f(complex) refers to the fraction of complex in each experiment, which was estimated using the $K_D$ value obtained from AlphaScreen experiments. In the case of the association reactions, f(complex) refers to the fraction of complex formed within the range of Rb concentrations used. In the case of the dissociation reactions, it refers to the fraction of complex displaced. # The second value corresponds to a second replicate done in another laboratory using a different batch of protein and a different stopped-flow apparatus.

**Table S4.** ***Effect of mutations in the TSE for binding of the LxCxE motif***

| *Mutant* | *Class of mutation* | *Class LEFFLER*# | *Class (Φ-values)*** |
|---|---|---|---|
| *D21A* | *side chain deletion* | *no interactions* | *no interactions* |
| *D21A neutral* | *dissection electrostatics* | *non-native energetics* | *ND* |
| *D21A electro* | *dissection electrostatics* | *native-like energetics* | *native-like energetics* |
| *L22A* | *side chain deletion* | *no interactions* | *no interactions* |
| *L22I* | *side chain replacement conservative* | *no interactions* | *no interactions* |
| *I22A* | *side chain deletion* | *no interactions* | *no interactions* |
| *Y23A* | *side chain deletion* | *no interactions* | *no interactions* |
| *Y23F* | *-OH deletion* | *no interactions* | *ND* |
| *F23A* | *side chain deletion* | *no interactions* | *no interactions* |
| *Y23F$_{F25}$* | *-OH deletion* | *non-native energetics* | *ND* |
| *C24A* | *side chain deletion* | *no interactions* | *no interactions* |
| *C24S* | *side chain replacement conservative* | *no interactions* | *no interactions* |
| *S24A* | *side chain deletion* | *no interactions* | *no interactions* |
| *Y25A* | *side chain deletion* | *no interactions* | *no interactions* |
| *Y25F* | *-OH deletion* | *no interactions* | *no interactions* |
| *Y25H* | *side chain replacement conservative* | *no interactions* | *no interactions* |
| *F25A* | *side chain deletion* | *no interactions* | *ND* |
| *H25A* | *side chain deletion* | *no interactions* | *no interactions* |
| *H25F* | *side chain replacement conservative* | *no interactions* | *ND* |
| *Y25F$_{F23}$* | *-OH deletion* | *non-native energetics* | *non-native energetics* |
| *Y23F-Y25F* | *-OH deletion* | *non-native energetics* | *non-native energetics* |
| *Y23F-Y25F neutral* | *dissection electrostatics* | *non-native energetics* | *non-native energetics* |
| *Y23F-Y25F electro* | *dissection electrostatics* | *native-like energetics* | *ND* |
| *E26A* | *side chain deletion* | *native-like energetics* | *native-like energetics* |
| *E26A neutral* | *dissection electrostatics* | *non-native energetics* | *non-native energetics* |
| *E26A electro* | *dissection electrostatics* | *native-like energetics* | *native-like energetics* |
| *E26D* | *side chain replacement conservative* | *native-like energetics* | *no interactions* |
| *E26D neutral* | *dissection electrostatics* | *no interactions* | *no interactions* |
| *E26D electro* | *dissection electrostatics* | *native-like energetics* | *native-like energetics* |
| *D26A* | *side chain deletion* | *native-like energetics* | *native-like energetics* |
| *D26A neutral* | *dissection electrostatics* | *native-like energetics* | *native-like energetics* |
| *D26A electro* | *dissection electrostatics* | *native-like energetics* | *native-like energetics* |
| *Q27A* | *side chain deletion* | *no interactions* | *ND* |
| *L28A* | *side chain deletion* | *native-like energetics* | *native-like energetics* |
| *L28+3* | *side chain replacement conservative* | *native-like energetics* | *native-like energetics* |
| *L28+4* | *side chain replacement conservative* | *native-like energetics* | *native-like energetics* |

### Native-like energetics if $\Delta\Delta G$ values equal sign and $|\Delta\Delta G| > 0.2$; Non-native energetics: $\Delta\Delta G^{\ddagger} < -0.2$ kcal/mol & 2*SD < 0.2 ** Native-like energetics if $\Phi > 0.2$; Non-native if $\Phi < 0$

**Table S5.** ***Experimental conditions used to determine the NaCl dependence of the equilibrium and rate constants for LxCxE:Rb interactions.***

| | ***Association reactions*** | | | | ***Dissociation reactions*** | | | |
|---|---|---|---|---|---|---|---|---|
| | **[Rb] (nM)** | **[FITC-peptide] (nM)** | **TB (s)*** | **traces** | **[Complex] (nM)** | **[peptide] (nM)** | **TB (s)*** | **traces** |
| **WT** | 500 | 100 | 1; 2 and 4 | > 9 | 250 | 5000 | 100 | > 5 |
| **D21A** | 500 | 100 | 1; 2 and 4 | > 8 | 1250 | 12500 | 10 | > 6 |
| **E26A** | 3000 | 400 | 0.2 | > 7 | 5000 | 50000 | 0.4 | 5 |
| **E26D** | 1000 | 200 | 1 and 2 | > 8 | 5000 | 50000 | 2 | 5 |
| **Y23F-Y25F** | 1000 | 100 | 0.4 and 1 | > 8 | 500 | 1000 | 6 and 10 | > 8 |

*TB refers to the time base used for the trace recording in the different experiments. In some cases more than one time base was used in order to capture the completion of the trace for every salt concentration.

**Table S6.** ***NaCl dependence of the equilibrium and rate constants for the*** **[LxCxE$_{WT}$:Rb]** ***complex***

| **[NaCl] (M)** | $k_{on}$ ($M^{-1}s^{-1}$) | $k_{off}$ ($s^{-1}$) | $K_D$ (M) | $\Delta G^0$ (kcal/mol) |
|---|---|---|---|---|
| 0.08 | 3.64 ± 0.18 x $10^7$<br>3.21 ± 0.17 x $10^7$ | 0.137 ± 0.001<br>0.115 ± 0.001 | 3.76 ± 0.19 x $10^{-9}$<br>3.59 ± 0.32 x $10^{-9}$ | -11.29 ± 0.03<br>-11.32 ± 0.03 |
| 0.13 | 2.46 ± 0.12 x $10^7$<br>2.19 ± 0.11 x $10^7$ | 0.143 ± 0.001<br>0.133 ± 0.001 | 5.79 ± 0.29 x $10^{-9}$<br>6.07 ± 0.31 x $10^{-9}$ | -11.04 ± 0.03<br>-11.01 ± 0.03 |
| 0.2 | 1.57 ± 0.08 x $10^7$<br>1.45 ± 0.07 x $10^7$ | 0.156 ± 0.001<br>0.129 ± 0.001 | 9.94 ± 0.49 x $10^{-9}$<br>8.87 ± 0.45 x $10^{-9}$ | -10.72 ± 0.03<br>-10.79 ± 0.03 |
| 0.2* | 1.95 ± 0.12 x $10^7$<br>1.76 ± 0.11 x $10^7$ | 0.139 ± 0.001<br>0.126 ± 0.001 | 7.14 ± 0.47 x $10^{-9}$<br>7.19 ± 0.45 x $10^{-9}$ | -10.93 ± 0.04<br>-10.92 ± 0.04 |
| 0.38 | 8.19 ± 0.41 x $10^6$<br>7.02 ± 0.36 x $10^6$ | 0.180 ± 0.001<br>0.144 ± 0.001 | 2.20 ± 0.11 x $10^{-8}$<br>2.04 ± 0.11 x $10^{-8}$ | -10.26 ± 0.03<br>-10.31 ± 0.03 |
| 0.48 | 6.91 ± 0.35 x $10^6$<br>5.73 ± 0.30 x $10^6$ | 0.193 ± 0.001<br>0.152 ± 0.001 | 2.80 ± 0.14 x $10^{-8}$<br>2.65 ± 0.14 x $10^{-8}$ | -10.12 ± 0.03<br>-10.15 ± 0.03 |
| 0.68 | 4.70 ± 0.24 x $10^6$<br>4.25 ± 0.22 x $10^6$ | 0.189 ± 0.001<br>0.186 ± 0.001 | 4.00 ± 0.20 x $10^{-8}$<br>4.37 ± 0.23 x $10^{-8}$ | -9.91 ± 0.03<br>-9.86 ± 0.03 |

The second value corresponds to the results for the second replicate done in another laboratory using a different batch of protein and a different stopped-flow apparatus. This second replicate tested higher salt concentrations (0.8 and 1 M). *Results for 200 mM NaCl determined as a function of Rb Concentration (PFO plots).

**Table S7.** ***NaCl dependence of the equilibrium and rate constants for the*** **[LxCxE$_{D21A}$:Rb]** ***complex***

| [NaCl] (M) | $k_{on}$ ($M^{-1}s^{-1}$) | $k_{off}$ ($s^{-1}$) | $K_D$ (M) | $\Delta G^0$ (kcal/mol) | $\Delta\Delta G^0$ (kcal/mol) | $\Delta\Delta G^{\ddagger}$ (kcal/mol) | ϕ-value |
|---|---|---|---|---|---|---|---|
| 0.08 | 2.84 ± 0.14 x $10^7$ | 1.451 ± 0.004 | 5.10 ± 0.26 x $10^{-8}$ | -9.77 ± 0.03 | 1.52 ± 0.04 | 0.14 ± 0.04 | 0.10 ± 0.03 |
| 0.13 | 2.23 ± 0.12 x $10^7$ | 1.524 ± 0.004 | 6.70 ± 0.39 x $10^{-8}$ | -9.61 ± 0.03 | 1.43 ± 0.04 | 0.05 ± 0.04 | 0.03 ± 0.03 |
| 0.2 | 1.61 ± 0.82 x $10^7$ | 1.622 ± 0.005 | 1.01 ± 0.51 x $10^{-7}$ | -9.38 ± 0.03 | 1.35 ± 0.04 | -0.02 ± 0.04 | -0.01 ± 0.03 |
| 0.2* | 1.75 ± 0.11 x $10^7$ | 1.37 ± 0.03 | 7.83 ± 0.50 x $10^{-8}$ | -9.53 ± 0.04 | 1.39 ± 0.05 | 0.06 ± 0.05 | 0.05 ± 0.04 |
| 0.38 | 1.02 ± 0.05 x $10^7$ | 1.591 ± 0.007 | 1.56 ± 0.08 x $10^{-7}$ | -9.12 ± 0.03 | 1.14 ± 0.04 | -0.13 ± 0.04 | -0.11 ± 0.04 |
| 0.48 | 8.97 ± 0.46 x $10^6$ | 1.642 ± 0.002 | 1.83 ± 0.09 x $10^{-7}$ | -9.03 ± 0.03 | 1.10 ± 0.04 | -0.15 ± 0.04 | -0.14 ± 0.04 |
| 0.68 | 7.43 ± 0.40 x $10^6$ | 1.295 ± 0.007 | 1.74 ± 0.09 x $10^{-7}$ | -9.06 ± 0.03 | 0.86 ± 0.04 | -0.27 ± 0.04 | -0.31 ± 0.05 |

*Results for 200 mM NaCl determined as a function of Rb Concentration (PFO plots)

**Table S8.** ***NaCl dependence of the equilibrium and rate constants for the*** **[LxCxE$_{E26D}$:Rb]** ***complex***

| [NaCl] (M) | $k_{on}$ ($M^{-1}s^{-1}$) | $k_{off}$ ($s^{-1}$) | $K_D$ (M) | $\Delta G^0$ (kcal/mol) | $\Delta\Delta G^0$ (kcal/mol) | $\Delta\Delta G^{\ddagger}$ (kcal/mol) | ϕ-value |
|---|---|---|---|---|---|---|---|
| 0.08 | 2.51 ± 0.26 x $10^7$ | 5.479 ± 0.008 | 2.18 ± 0.23 x $10^{-7}$ | -8.93 ± 0.06 | 2.36 ± 0.07 | 0.22 ± 0.07 | 0.09 ± 0.03 |
| 0.13 | 1.48 ± 0.13 x $10^7$ | 6.440 ± 0.008 | 4.35 ± 0.38 x $10^{-7}$ | -8.53 ± 0.05 | 2.51 ± 0.06 | 0.30 ± 0.06 | 0.12 ± 0.02 |
| 0.2 | 1.03 ± 0.08 x $10^7$ | 6.585 ± 0.008 | 6.41 ± 0.51 x $10^{-7}$ | -8.30 ± 0.05 | 2.42 ± 0.06 | 0.25 ± 0.06 | 0.10 ± 0.02 |
| 0.2* | 1.04 ± 0.07 x $10^7$ | 5.446 ± 0.006 | 5.26 ± 0.33 x $10^{-7}$ | -8.42 ± 0.04 | 2.50 ± 0.05 | 0.37 ± 0.05 | 0.15 ± 0.02 |
| 0.38 | 5.10 ± 0.32 x $10^6$ | 7.245 ± 0.009 | 1.42 ± 0.09 x $10^{-6}$ | -7.84 ± 0.04 | 2.43 ± 0.05 | 0.28 ± 0.05 | 0.11 ± 0.02 |
| 0.48 | 5.07 ± 0.31 x $10^6$ | 7.256 ± 0.009 | 1. 43 ± 0.09 x $10^{-6}$ | -7.83 ± 0.04 | 2.29 ± 0.05 | 0.18 ± 0.05 | 0.08 ± 0.02 |
| 0.68 | 3.93 ± 0.23 x $10^6$ | 7.248 ± 0.009 | 1.85 ± 0.11 x $10^{-6}$ | -7.68 ± 0.03 | 2.23 ± 0.05 | 0.11 ± 0.04 | 0.05 ± 0.02 |

*Results for 200 mM NaCl determined as a function of Rb Concentration (PFO plots)

**Table S9.** ***NaCl dependence of the equilibrium and rate constants for the*** **[LxCxE$_{E26A}$:Rb]** ***complex***

| [NaCl] (M) | $k_{on}$ ($M^{-1}s^{-1}$) | $k_{off}$ ($s^{-1}$) | $K_D$ (M) | $\Delta G^0$ (kcal/mol) | $\Delta\Delta G^0$ (kcal/mol) | $\Delta\Delta G^{\ddagger}$ (kcal/mol) | ϕ-value |
|---|---|---|---|---|---|---|---|
| 0.08 | 4.80 ± 0.43 x $10^6$ | 52.6 ± 0.6 | 1.10 ± 0.10 x $10^{-5}$ | -6.65 ± 0.05 | 4.64 ± 0.06 | 1.18 ± 0.06 | 0.254 ± 0.01 |
| 0.13 | 3.96 ± 0.34 x $10^6$ | 46.0 ± 0.5 | 1.16 ± 0.10 x $10^{-5}$ | -6.61 ± 0.05 | 4.42 ± 0.06 | 1.06 ± 0.06 | 0.240 ± 0.01 |
| 0.2 | 3.50 ± 0.28 x $10^6$ | 36.9 ± 0.3 | 1.05 ± 0.09 x $10^{-5}$ | -6.67 ± 0.05 | 4.05 ± 0.06 | 0.87 ± 0.06 | 0.215 ± 0.01 |
| 0.2* | 3.4 ± 0.4 x $10^6$ | 37.76 ± 0.3 | 1.10 ± 0.1 x $10^{-5}$ | -6.65 ± 0.06 | 4.27 ± 0.07 | 1.01 ± 0.07 | 0.24 ± 0.02 |
| 0.68 | 1.97 ± 0.14 x $10^5$ | 23.2 ± 0.1 | 1.17 ± 0.08 x $10^{-5}$ | -6.61 ± 0.04 | 3.30 ± 0.05 | 0.50 ± 0.05 | 0.153 ± 0.02 |

*Results for 200 mM NaCl determined as a function of Rb Concentration (PFO plots).

**Table S10.** ***NaCl dependence of the equilibrium and rate constants for the*** **[LxCxE$_{Y23F-Y25F}$:Rb]** ***complex***

| [NaCl] (M) | $k_{on}$ ($M^{-1}s^{-1}$) | $k_{off}$ ($s^{-1}$) | $K_D$ (M) | $\Delta G^0$ (kcal/mol) | $\Delta\Delta G^0$ (kcal/mol) | $\Delta\Delta G^{\ddagger}$ (kcal/mol) | ϕ-value |
|---|---|---|---|---|---|---|---|
| 0.08 | 3.99 ± 0.21 x $10^7$ | 1.43 ± 0.004 | 3.59 ± 0.05 x $10^{-8}$ | -9.98 ± 0.03 | 1.34 ± 0.04 | -0.13 ± 0.04 | -0.10 ± 0.03 |
| 0.13 | 2.85 ± 0.15 x $10^7$ | 1.58 ± 0.006 | 5.55 ± 0.09 x $10^{-8}$ | -9.72 ± 0.03 | 1.29 ± 0.04 | -0.15 ± 0.04 | -0.12 ± 0.03 |
| 0.2 | 1.89 ± 0.10 x $10^7$ | 1.76 ± 0.008 | 9.33 ± 0.14 x $10^{-8}$ | -9.42 ± 0.03 | 1.37 ± 0.04 | -0.15 ± 0.04 | -0.11 ± 0.03 |
| 0.2* | 2.1 ± 0.2 x $10^7$ | 1.68 ± 0.003 | 8.15 ± 0.8 x $10^{-8}$ | -9.51 ± 0.06 | 1.41 ± 0.05 | -0.03 ± 0.07 | -0.02 ± 0.05 |
| 0.38 | 1.01 ± 0.06 x $10^7$ | 2.00 ± 0.12 | 1.97 ± 0.17 x $10^{-7}$ | -8.99 ± 0.05 | 1.32 ± 0.06 | -0.21 ± 0.04 | -0.16 ± 0.03 |
| 0.48 | 8.58 ± 0.47 x $10^6$ | 2.01 ± 0.02 | 2.34 ± 0.05 x $10^{-7}$ | -8.89 ± 0.03 | 1.27 ± 0.04 | -0.24 ± 0.04 | -0.19 ± 0.04 |
| 0.68 | 6.24 ± 0.37 x $10^6$ | 2.03 ± 0.02 | 3.25 ± 0.09 x $10^{-7}$ | -8.69 ± 0.04 | 1.17 ± 0.05 | -0.22 ± 0.05 | -0.19 ± 0.04 |

*Results for 200 mM NaCl determined as a function of Rb Concentration (PFO plots).

**Table S11.** ***Analysis of the dependence of association and dissociation rate constants on [NaCl] according to the Debye-Hückel theory.***

| | $U_{on}$ (kcal/mol) | $U_{off}$ (kcal/mol) | $k_{on,\ basal}$ | $k_{off,\ basal}$ | $\Delta\Delta G^{0}_{N}$ (kcal/mol) | $\Delta\Delta G^{0}_{E}$ (kcal/mol) | $\Delta\Delta G^{\ddagger}_{N}$ (kcal/mol) | $\Delta\Delta G^{\ddagger}_{E}$ (kcal/mol) | $\Phi_N$ | $\Phi_E$ | *a* (Å)§ |
|---|---|---|---|---|---|---|---|---|---|---|---|
| **WT** | -5.4 ± 0.2<br>-5.5 ± 0.2 | 1.14 ± 0.01<br>0.94 ± 0.01 | (1.44 ± 0.2)x10$^5$<br>(1.20 ± 0.2)x10$^5$ | 0.422 ± 0.003<br>0.306 ± 0.001 | -<br>- | -<br>- | -<br>- | -<br>- | -<br>- | -<br>- | |
| **D21A** | -3.7 ± 0.2 | 0.10 ± 0.01 | (6.64 ± 0.9)x10$^6$ | 1.67 ± 0.02 | -0.09 ± 0.03 | 2.75 ± 0.11 | -0.89 ± 0.02 | 1.72 ± 0.05 | -† | 0.63 ± 0.05 | |
| **E26D** | -4.7 ± 0.2 | 0.64 ± 0.01 | (1.77 ± 0.3)x10$^5$ | 11.42 ± 0.04 | 1.80 ± 0.16 | 1.21 ± 0.08 | -0.12 ± 0.02 | 0.72 ± 0.05 | -0.066 ± 0.18 | 0.60 ± 0.11 | 6 |
| **E26A** | -2.3 ± 0.3 | -2.17 ± 0.03 | (4.4 ± 1.0)x10$^5$ | 5.72 ± 0.12 | 0.86 ± 0.03 | 6.37 ± 0.04 | -0.65 ± 0.02 | 3.06 ± 0.11 | -0.75 ± 0.05 | 0.48 ± 0.04 | |
| **D26A** | 2.34 ± 0.4 | -2.81 ± 0.03 | (3.6 ± 0.7)x10$^5$ | 0.212 ± 0.03 | -0.94 ± 0.16 | 5.16 ± 0.09 | -0.53 ± 0.03 | 2.34 ± 0.13 | 0.6 ± 0.2 | 0.46 ± 0.06 | |
| **Y23F-Y25F** | -5.0 ± 0.2 | 1.10 ± 0.02 | (2.5 ± 0.4)x10$^6$ | 4.44 ± 0.09 | 1.14 ± 0.04 | 0.34 ± 0.16 | -0.42 ± 0.02 | 0.50 ± 0.04 | -0.37 ± 0.05 | -† | |

The NaCl dependence of the kinetic association and dissociation rate constants was globally fitted for all mutants using Eq. (8) from Materials and Methods (SI Appendix, Figure S8). The values of ln $k_{on,\ basal}$ and $U_{on}$/RT or those of ln $k_{off,\ basal}$ and $U_{off}$/RT were obtained as fitting parameters for each mutant. The constant *b* was kept as a global fitting parameter and *a* was calculated as *a* = *b/b'*(See Eq. (6-8) in Materials and Methods).

†The $\Phi_{E/N}$ for is not reported because $\Delta\Delta G^{0}_{E/N}$ < 0.6 kcal/mol.

§The parameter was fixed to 6Å when fitting our data as described in Materials and Methods.

*The second value for WT corresponds to a second replicate done in another laboratory using a different batch of protein and a different stopped-flow apparatus.

**Table S12.** ***Secondary structure analysis of the LxCxE peptide by NMR and Far-UV CD.***

| | NMR# | | | | | | | Far-UV CD* | |
|---|---|---|---|---|---|---|---|---|---|
| | δ2Δ method | | | | CSI | | | helicity (%) | |
| | helicity (%) | polypoline II (PPII) (%) | Coil (%) | Extended beta (%) | helicity (%) | Coil (%) | Extended beta (%) | Method 1 | Method 2 |
| **WT** | 0.2 | 17.7 | 80.2 | 2.0 | 5.3 | 70.5 | 24.4 | 5.0 | 8.6 |
| **Y23F** | 0.7 | 18.3 | 79.0 | 2.1 | 7.5 | 66.9 | 25.8 | 4.6 | 8.2 |
| **Y25F** | 0.3 | 17.5 | 81.0 | 1.2 | 6.4 | 68.5 | 25.0 | 5.3 | 8.8 |
| **Y23F-Y25F** | 0.9 | 18.3 | 79.3 | 1.6 | 7.8 | 67.2 | 25.1 | 6.6 | 10.2 |

#Secondary structure analysis by NMR of LxCxE peptide mutants with NMR data processed by the δ2Δ method or by secondary chemical shifts (CSI) (See Material and Methods for more information). For both, helical fraction = 1 corresponds to 100% helicity.

* Helicity prediction by CD. This can be calculated by using two different methods (see Material and Methods for more information). Method 1 uses Eq. (20) from [1] and Method 2 uses Eq. (21) from [2] (See material and methods for more information).

#### Supplementary Text

##### Ground state effects

Most IDRs studied to date present variable (10-50 %) helical propensity in the unbound state[3–9], and acquire helical structure upon binding to disordered[7,10,11] or folded[5–7,12–15] partners, but ground state controls are only available for a handful of systems[5,6,10,15,16]. Ground state effects can confound the assignment of native-like or non-native energetics in the TSE from Φ-value or Leffler plot analysis. For example, apparent native-like energetics in the TSE for folding of the NTL9 protein originate from oppositely charged residues that establish non-native interactions in the denatured state, which persist in the TSE[17]. For a few IDRs[12,13] Φ-value analysis revealed high ($\alpha \sim 0.8$) LFER values that suggested a degree of structuring higher than the global average of protein folding reactions ($\alpha \sim 0.3$)[18,19]. We suggest that yet-unprobed non-native electrostatic interactions present in the ground state of these highly charged IDRs may explain this unusual energetics, much like the case of NTL9[17,20]. This emphasizes that while IDR binding mechanisms may be diverse, ground state controls are essential to achieve a precise picture of the TSE energetics.

##### Far-UV Circular Dichroism (Far-UV CD) analysis of LxCxE peptides

All peptides have a Far-UV CD spectrum (SI Appendix, Figure S10) with a minimum around 198 nm and a small shoulder around 220 nm, which are characteristic of disordered polypeptides[21–23]. Most spectra from mutant peptides show an excellent overlap with the $LxCxE_{WT}$ peptide with changes occurring upon mutation of aromatic residues at positions 23 and 25 being explained by the known contribution of aromatic side chains in the far-UV region of peptide and protein CD spectra[24,25]. Specifically, two positive contributions arise from the aromatic side chains[26,27]: one around 200 nm corresponding to the amide π-π* transition and another between 220-230 nm corresponding to the n-π* transition. In Y23A and Y25A, where only one aromatic residue is removed, the positive contribution at 200 nm becomes smaller, resulting in an increased negative signal at both 200 nm and 230 nm. This effect is more pronounced when both aromatic residues are removed, as observed in the Y23A-Y25A mutant. For the Y-to-F mutation, there is a change in the identity of the aromatic residue. The electron transitions of phenylalanine have a lower intensity and occur at a higher energy than those of tyrosine[28], which explains the shift of the positive band to a lower wavelength. It is interesting to note that mutations at position 23 do not have the same effect as those at position 25 (e.g. Y23A vs Y25A). In addition, the changes in the spectra of the double mutants (Y23A-Y25A and Y23F-Y25F) do not result from the sum of the effects observed with individual mutations. This discrepancy arises from the known non-additive contributions of aromatic residue transitions [29]. For L28+4, the shift in the position of the minimum from 198 nm to 200 nm and a slight increase in signal at 220 nm may reflect a very marginal increase in fractional helical populations due to the addition of two alanine residues to introduce the "spacer" mutation, but the spectrum indicates that L28+4 remains largely disordered.

##### Change in hydrophobicity

Non-native energetics could arise if a hydrophobic moiety stabilizes the TSE by improving the interactions with other loosely packed hydrophobic residues but destabilizes the bound state due to poor fitting with the remaining tightly packed hydrophobic core[30]. For some binding reactions involving IDPs nonspecific hydrophobic interactions play a key role in stabilizing the TSE for binding[31–33] and a signature of this phenomenon is a high correlation between the change in hydrophobicity measured as Δlog P and the $\Delta\Delta G^{\ddagger}$ or $\Delta\Delta G^{0}$ values. For example for the APP:PARM complex[33] the high $R^2$ values and slope for Δlog P versus $\Delta\Delta G^{\ddagger}$ **(**$R^2$ = 0.7, slope = -3.4**)** and $\Delta\Delta G^{0}$ ($R^2$ = 0.8, slope = -1.0) indicate that hydrophobic interactions stabilize both the TSE and the complex at equilibrium. In contrast, other IDP-domain interactions[14,34–37] show a lack of correlation between Δlog P and $\Delta\Delta G^{\ddagger}$ ($R^2$ = 0.006-0.40) [33] indicating that hydrophobic interactions have a negligible contribution to the TSE for binding[32]. To determine whether nonspecific hydrophobic interactions played a role in the TSE for $LxCxE_{WT}$ binding to Rb, we calculated the change in the hydrophobicity produced by each mutation, expressed as *Δlog* P (See Materials and Methods), and assessed its correlation with $\Delta\Delta G^{\ddagger}$ and $\Delta\Delta G^{0}$ (SI Appendix, Table S2). The $LxCxE_{WT}$ mutations span a very similar range in *Δlog* P, $\Delta\Delta G^{\ddagger}$ or $\Delta\Delta G^{0}$ as the above-mentioned studies. The plot of *Δlog* P versus $\Delta\Delta G^{\ddagger}$ or $\Delta\Delta G^{0}$ for $LxCxE_{WT}$ binding to Rb (SI Appendix, Figure S12 A,B) yielded low values for the slopes (-0.21 for $\Delta\Delta G^{\ddagger}$ and -0.12 for $\Delta\Delta G^{0}$) and low $R^2$ values (0.003 for $\Delta\Delta G^{\ddagger}$ and 0.013 for $\Delta\Delta G^{0}$) indicating no correlation between changes in the free energy of binding and changes in hydrophobicity, both for the TSE ($\Delta\Delta G^{\ddagger}$) and at equilibrium ($\Delta\Delta G^{0}$). This conclusion remains unchanged when considering $\Delta\Delta G^{\ddagger}$ values derived from the non-electrostatic component for D21A, E26A, E26D, D26A, and Y23F–Y25F, which show weak correlations (slope = −1.8, $R^2$ = 0.22 for $\Delta\Delta G^{\ddagger}$; slope = −0.40, $R^2$ = 0.13 for $\Delta\Delta G^{0}$; SI Appendix, Figure S12C and D). These results favor the interpretation that nonspecific hydrophobic interactions are unlikely to play a major role in stabilizing the TSE for binding of $LxCxE_{WT}$ to Rb and instead suggest that the non-native energetics originate from non-favorable contacts established by moieties of D21, Y23, Y25 and E26 that introduce energetic frustration in the TSE for binding.
